# Intrinsically Disordered Protein Coating for Oral Delivery of Peptide Drugs

**DOI:** 10.1101/2025.10.23.684059

**Authors:** Max Ney, Parul Sirohi, Yulia Shmidov, Anurag Singh, Gable Wadsworth, Xinghai Li, James Zheng, Erica Peng, Lixin Fan, Tharun Selvam Mahendran, Sonal Deshpande, Navya Tripathi, Jonathan C. Su, Joshua James Milligan, Yun-Xing Wang, Priya R. Banerjee, Ashutosh Chilkoti

## Abstract

Advancing oral delivery of peptide therapeutics requires innovative materials that overcome gastrointestinal barriers. We introduce the first engineered synthetic intrinsically disordered protein (SynIDP) that self-assembles into an enteric coating, encapsulating peptide drugs to enhance gastric acid resistance and intestinal targeting. This SynIDP recapitulates the molecular design principles and phase transitions of native IDPs to exhibit temperature-controlled condensation and pH-controlled solidification—both transitions being reversible and precisely tuned to intestinal cues. Through detailed analysis of the kinetics of the liquid-to-solid phase transition, we achieve control over the nano-to-microscale morphology of the protein coating, optimizing drug encapsulation and protection. The coating protects peptide-based weight loss drugs for over 60 minutes in simulated gastric conditions, then dissolves to release the active compound. Oral delivery to obese mice results in more consistent weight loss compared to the unencapsulated drug. This modular protein-based coating is a promising platform technology for enhancing oral peptide drug delivery and improving patient compliance.

## Main

Oral delivery of peptide therapeutics remains a major unmet need in metabolic and other diseases. In the past decade, glucagon-like peptide-1 receptor agonists (GLP1-RAs) such as semaglutide, liraglutide, and exenatide have transformed the treatment of diabetes and obesity by modulating blood glucose and appetite [1, 2]. While co-formulation with permeation enhancers like salcaprozate sodium (SNAC) [3] has enabled the first FDA-approved oral GLP1-RA for obesity, most peptide drugs—including those for cancer, osteoporosis, HIV, and infectious diseases [4] — still require injection due to poor oral bioavailability and the need for fasting to reduce gastric acid volume. These barriers limit patient adherence and restrict the therapeutic potential of peptide drugs.

Here, we report a protein-based enteric coating inspired by biomolecular condensates that enables the oral delivery of GLP1-RAs without the need for fasting. Our approach draws on the phase separation principles that underlie condensates of naturally occurring intrinsically disordered proteins (IDPs) [5], but encodes them directly into a genetically engineered synthetic variant— that we term SynIDP—that is designed at the sequence level to integrate drug-specific supramolecular binding, pH-responsive reversible self-assembly, and resistance to gastric degradation into a modular platform.

Our design of the SynIDP combines temperature-dependent liquid-liquid phase separation (LLPS) with pH-triggered solidification, encoded in distinct multivalent domains. To demonstrate the utility of this platform, we encapsulate GLP1-RA peptide drugs using two strategies: 1) supramolecular co-assembly by genetic fusion to an amyloid fiber-forming domain, and 2) nonspecific hydrophobic interactions with the alkyl chain of semaglutide.

In vitro, dynamically arrested spherical condensates of the SynIDP with a high-volume and low surface area maximize drug loading and survival in simulated gastric acid (SGA), retaining 40% of undigested GLP1-RA activity upon release. In vivo studies in fed obese mice show that the encapsulated oral formulation achieves weight loss comparable to subcutaneously injected drug, whereas unencapsulated oral drug does not. Imaging and pharmacokinetic analyses reveal that active drug dissociates from the coating in the upper intestine and is absorbed in the lower intestine.

This work establishes engineered SynIDP condensates as a versatile enteric delivery platform for peptide therapeutics. The modularity of the self-assembly domains, combined with precise control over condensate formation and material properties, enables adaptation to a wide range of acid-labile drugs. and opens new avenues for oral delivery of biologics in metabolic disease and beyond.

## Inspiration from Natural Condensates

Biomolecular condensates are membraneless organelles that in many instances are formed through LLPS of native intrinsically disordered proteins (IDPs) and play key roles in numerous cellular processes [6]. Many well-studied condensate-forming proteins—such as FUS [7], hnRNPA1 [8], and TDP-43 [9]—contain intrinsically disordered regions (IDRs) interspersed with multiple short, fiber-forming peptides known as reversible amyloid cores (RACs) (**Fig. 1a**) [10]. IDRs lack stable secondary structures and promote liquid-like condensate properties through multivalent interactions [5], whereas RACs form strong self-interactions that drive fibril or aggregate formation [11]. Together, these self-assembly modes can cause condensates to gradually undergo dynamical arrest or transition to amyloid fibrils in a process commonly known as condensate aging [12–14]. The interplay between fiber assembly and LLPS is relevant to both degenerative diseases and normal cellular functions; however, few systems to date have harnessed these principles for the design of functional biomaterials, as we do here.

**Fig. 1:**
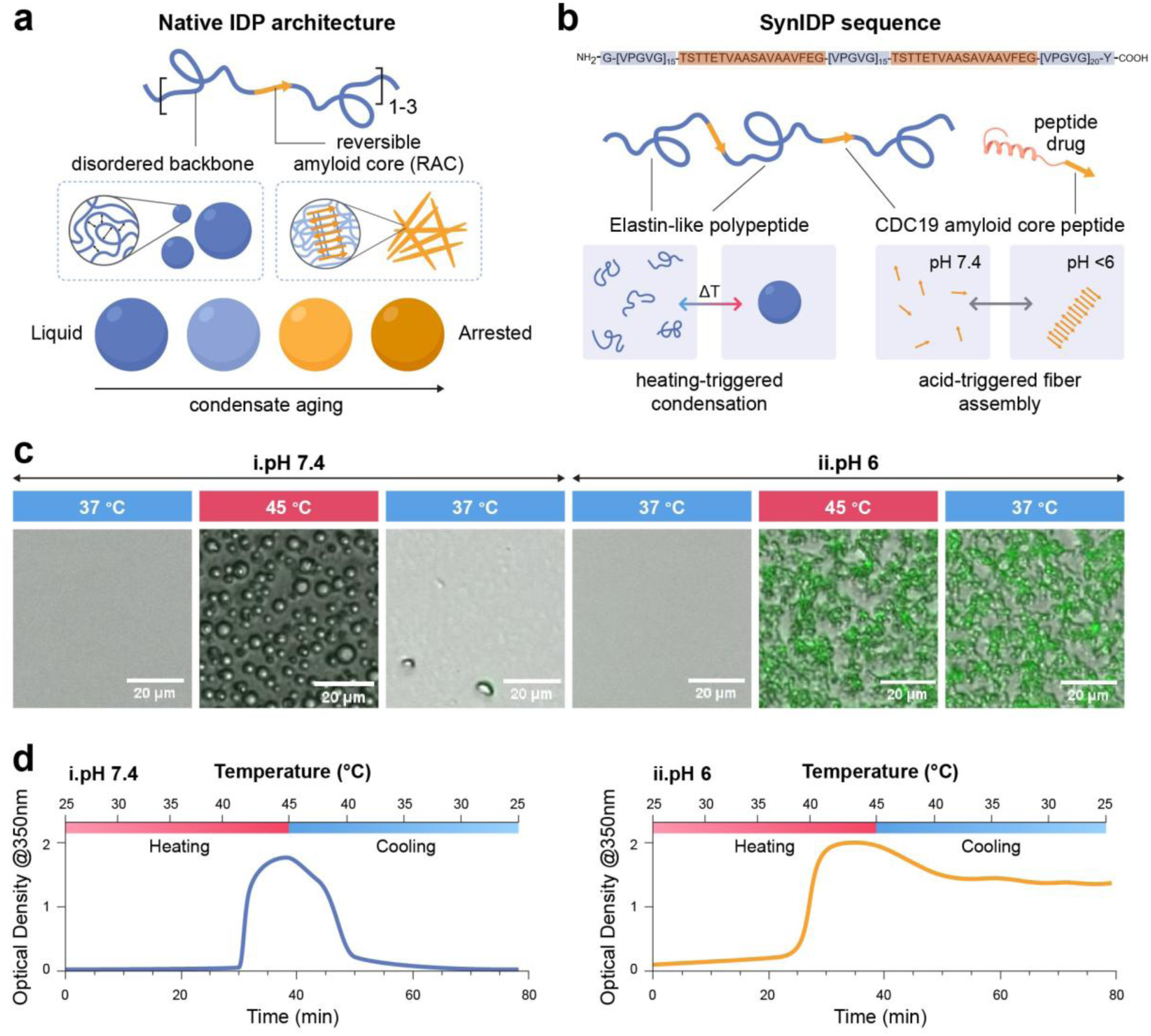
Design of a synIDP with GI-tailored phase transitions. **a**, Native IDPs are primarily composed of intrinsically disordered peptides but can also contain 1-3 fiber-forming peptides called reversible amyloid cores (RACs) which cause condensates to age into arrested states or fiber plaques. **b,** The molecular design of the synIDP for enteric drug delivery (top). The synIDP contains three temperature-responsive ELP tracts which reversibly drive synIDP condensation, and two pH-responsive RAC peptides to drive aging and co-assembly with a peptide drug that contains one copy of the RAC domain. **c,** Bright field and amyloid dye fluorescence microscopy of the synIDP in PBS at pH 7.4 (**i)** and pH 6 (**ii**) demonstrate thermal reversibility, amyloid dye intensity, and varied condensate morphologies. **d,** UV-Vis spectroscopy of heating and cooling at pH 7.4 (**i**) or pH 6 (**ii**) in PBS characterize the transition above body temperature and reversible-irreversible switch with changing pH.

## SynIDP Design

We engineered a synthetic intrinsically disordered protein (SynIDP) with temperature– and pH-responsive condensation and RAC-mediated fiber formation (**Fig. 1a**). The condensation domain is a temperature-sensitive elastin-like polypeptide (ELP) that undergoes thermally reversible LLPS at a transition temperature (T_t_) that is tunable by its sequence [15], chain length [16], salt [17], and fusion partners [18], and their structural disorder promotes condensate formation without fibril-driven aging (**Fig. 1b**). The ELP is composed of 50 repeats of VPGVG motifs from human tropoelastin [19], chosen to approximate the length of native IDRs [7, 9, 12] and to exclude aromatic residues, thereby limiting pepsin-mediated cleavage in the stomach [20]. Fibril formation was programmed by embedding two copies of the pH-responsive RAC from yeast kinase CDC19—after the 15^th^ and 30^th^ repeats of the ELP—that is known to assemble into nanofibrils below pH 5.7 and to protect CDC19 from proteolysis during stress (**Fig. 1b**)[21]. SynIDPs were expressed in *E. coli* and purified by inverse transition cycling (ITC), a non-chromatographic method that exploits the LLPS behavior of ELPs to centrifugally clarify them from non-thermally responsive contaminants [18].

## Phase Separation is Triggered by Temperature and Liquid-to-Solid Transition is Controlled by pH

Overlay of brightfield images and fluorescence images utilizing an amyloid reporter dye, Thioflavin T (ThT), revealed that at pH 7.4, the protein solution remained clear and ThT-negative at 37°C. Upon heating to 45°C, the protein solution underwent temperature-triggered LLPS into droplets (**Fig. 1c(i)**). These droplets remained ThT-negative, suggesting the formation of a condensate with liquid-like rather than amyloid properties. The droplets dissolved and reverted to a clear soluble phase upon cooling, confirming that phase separation is reversible.

At pH 6, heating induced the formation of arrested droplet networks that were ThT-positive, indicating a change in the internal structure of the droplets that persisted after cooling (**Fig. 1c(ii)**). UV–Vis spectroscopy (350 nm, 0.3°C/min) showed a reversible turbidity transition at 43 °C at pH 7.4 (**Fig. 1d(i)**) but an irreversible transition at 40 °C at pH 6 (**Fig. 1d(ii)**), indicating physical crosslinking by RAC motifs.

We investigated whether thermal reversibility could be dynamically switched in response to changing pH after the thermal transition. We heated samples at either pH 4 or 7 in citrate-phosphate buffer to access physiologically relevant pHs, then added acid or base to adjust the pH before cooling (**Supplementary** Fig. 1). The pH 4 condensate sample that was adjusted to pH 7 during cooling became thermally reversible, while the sample adjusted from pH 7 to 4 became thermally irreversible. These pH-switching experiments confirmed that condensate metastability is pH-dependent: condensates at pH 7 are reversible, whereas acidic pH locks condensates into a dynamically arrested state.

We hypothesized that the thermal irreversibility at acidic pH arises from a liquid-to-solid transition, likely driven by the glutamic acid residues in the RAC [21]. To test this hypothesis, we performed Fluorescence Recovery After Photobleaching (FRAP) to quantify protein mobility in condensates. Confocal images of Alexa Fluor 488–labeled SynIDP condensates at pH 6 or pH 7.4, 1 h after formation, reveal rapid recovery at neutral pH, as the bleached region disappears within 40 s (**Fig. 2a**). In contrast, condensates at pH 6 showed no visible fluorescence recovery after photobleaching. Quantification of the FRAP data, shown by the averaged recovery curves confirms ∼70% recovery at pH 7.4 versus none at pH 6 (**Fig. 2b**), indicating rapid exchange dynamics at neutral pH and dynamic arrest of the condensate under acidic conditions.

**Fig. 2:**
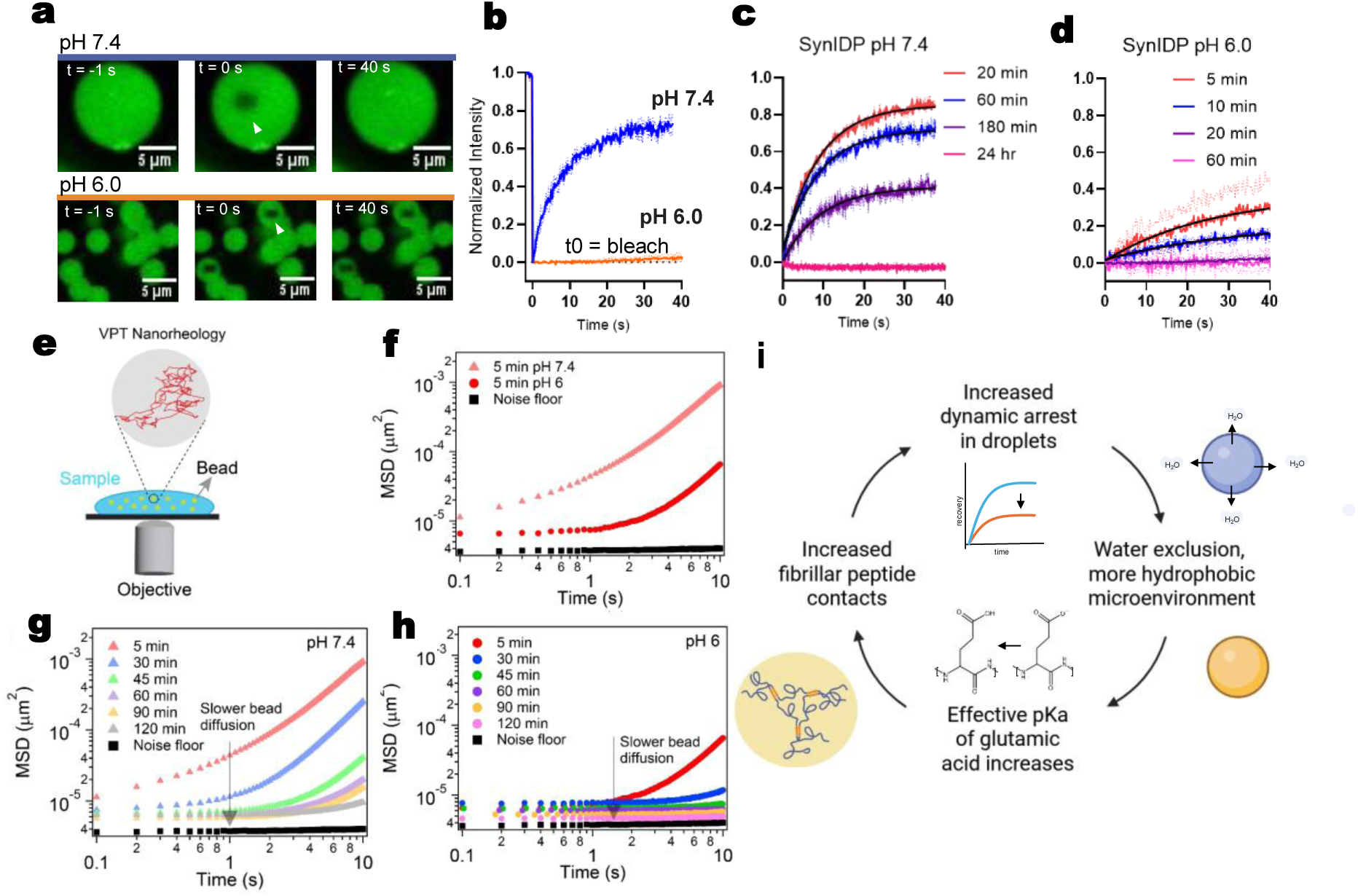
pH regulation of condensate aging stems from glutamines in the CDC19 RAC. **a**, Confocal images of Alexafluor-488 labeled synIDP condensates formed at pH 6 and 7.4 in PBS, which were photobleached and imaged to monitor their internal dynamics. **b,** The averaged intensities of three bleached regions were plotted over 40 seconds to compare droplets formed at pH 7.4 and pH 6.0. **c,** Recovery profiles at different droplet ages are shown for synIDP at pH 7.4 synIDP, and **d,** pH 6.0 synIDP. **e,** Schematic representation of video particle tracking (VPT) nanorheology used to estimate changes in the material properties of the synIDP condensates. **f,** Comparison of bead trajectories at two different pHs. Ensemble-averaged mean-squared displacement (MSD) of 200 nm beads embedded inside the condensates at pH 7.4 (pink, triangles) and pH 6 (red, circles). The detection limit is shown as the noise floor (black, squares). **g, h,** Age-dependent changes in the material properties of synIDP condensates at (f) pH 7.4 and (g) pH 6 are indicated by a downward shift in the MSDs of the beads. All measurements have been replicated independently at least n = 3 times. **i,** The mechanism for time and pH-dependent arrest is proposed to be from glutamine residues regulating aging. A positive feedback loop occurs in which dynamic arrest

We expected SynIDP droplets to age with time, with recovery decreasing as RAC interactions become increasingly dominant. Control FRAP curves for (VPGVG)₃₅ condensates at 60 and 180 min showed identical, fluid-like recovery (**Supplementary** Fig. 2). In contrast, SynIDP condensates at pH 7.4 progressively aged, with recovery falling to ∼40% at 180 min and complete arrest by 24 h (**Fig. 2d**). This multi-hour time scale of arrest parallels that of native IDP condensates of FUS under similar in vitro conditions [22].

At pH 6, aging was rapid: recovery was <40% at 5 min—the earliest measurable timepoint—and complete arrest occurred by 20 min as seen by the complete lack of recovery of the FRAP curves (**Fig. 2d**). We explored aging at intermediate pHs and observed FRAP recovery values that progressively decrease with both time and lower pH (**Supplementary Table 2 and Supplementary** Fig. 2), indicating that aging condensates access a spectrum of heterogenous states while undergoing dynamical arrest. Interestingly, the half-life of recovery only increases slightly before arrest – indicated as Tau values in the exponential fit of recovery curves – while fraction of recovery decreases dramatically (**Supplementary Table 3**). Dynamic acidification of 1 hour-old pH 7.4 condensates caused full arrest within 1 min at pH 5 (**Supplementary** Fig. 3 **left**). The reverse process, triggered by adding base to arrested pH 6 condensates, did not restore fluidity (**Supplementary** Fig. 3**, right)**, suggesting that dissolution at neutral or alkaline pH is not driven by bulk re-fluidization and likely occurs only at interfaces. Together, these results show that SynIDP condensate aging is pH-dependent, is driven solely by RAC peptides, and can be rapidly and dynamically tuned by environmental conditions.

Based upon FRAP data (**Fig. 2a-d**) and qualitative observations of distinct synIDP condensate dynamics under different solution conditions, we hypothesized that condensate dynamical arrest is mediated by a pH-dependent liquid-to-solid transition. To quantify this, we used video particle tracking (VPT) nanorheology [23, 24] (**Fig. 2e-h)**). This involves monitoring the thermal fluctuations of 200 nm diameter beads inside SynIDP condensates and estimating their mean-squared displacements (MSDs) (**Fig. 2e**). We observe that the ensemble-averaged MSDs of the embedded beads are generally higher at pH 7.4 (**Fig. 2f**) than at pH 6 (**Fig. 2f**) indicating that SynIDP condensates at pH 6 are more viscoelastic at the same condensate age. Additionally, we note that the bead motion in both conditions is sub-diffusive, displaying bending or flattening of the MSD trajectories and characterized by the diffusivity exponent α being less than 1[25, 26]. Both pH conditions show a shift towards lower MSD values over time (**Fig. 2g, h**), which is a hallmark of sample aging [25, 26]. Importantly, flattened MSDs were observed much earlier (after 45 minutes of sample age) in the case of pH 6 (**Fig. 2f**), compared to pH 7.4 which did not flatten over 120 minutes (**Fig. 2f**), confirming pH-dependent dynamical arrest of SynIDP condensates.

We hypothesized that condensates that are quenched deeper, i.e. a condition further from the binodal, into the two-phase regime would impact the material properties of condensates [27]. In SynIDP condensates, increasing the (NH_4_)_2_SO_4_ concentration to 500mM results in condensates that are more arrested and display enhanced sub-diffusivity than at 250 mM (**Supplementary** Fig. 4a). For comparison, viscoelastic ELP condensates exhibit an ∼11.5-fold decrease (from 23.62×10^-5^ to 2.07×10^-5^ µm^2^) in MSD when compared at the same salt concentrations (**Supplementary** Fig. 4a). ELP MSDs exhibited α = 1, allowing for the quantification of condensate viscosity, which demonstrate a 56-fold increase (from 3.07 ± 0.39 Pa.s to 122.01 ± 27.41 Pa.s) with the doubling of salt concentration (**Supplementary** Fig. 4a, b). This suggests that the dynamics of SynIDP condensates are dependent on quenching in addition to aging effects.

## Molecular Basis of pH-Responsive Arrest in SynIDP Condensates

Next, we investigated the molecular mechanism underlying pH-dependent solidification in SynIDP condensates, hypothesizing that protonation of two glutamic acid residues within each RAC domain reduces hydrophilicity and drives arrest. To test this hypothesis, we generated SynIDP variants in which each glutamic acid was independently substituted with alanine. Both substitutions led to markedly reduced protein yields, likely due to aggregation-induced insolubility. The double mutant could not be purified (**Supplementary** Fig. 5), consistent with severe aggregation of this SynIDP variant within the cell.

Strikingly, single substitutions abolish pH sensitivity, yielding irreversible transitions across all tested pH values, including pH 8 (**Supplementary** Fig. 6). These results demonstrate that both glutamic acids are essential for reversible, pH-tunable phase behavior and aging. The N-terminal substitution exhibits a sigmoidal phase separation profile (**Supplementary** Fig. 6**, left**), whereas the C-terminal mutation—adjacent to the sole aromatic residue—has a linear aggregation transition (**Supplementary** Fig. 6**, right**). This suggests that the C-terminal glutamic acid modulates aggregation via its proximity to a phenylalanine in the RAC, implicating aromatic stacking interactions that are known to stabilize amyloid fibrils[28].

Although the intrinsic pKa of glutamic acid is ∼4.25, hydrophobic environments can elevate this value above 6 [29]. Notably, Urry *et al.* reported a pKa of 6.1 for a glutamic acid within a phase-separated ELP 10-mer in saline [30]. We propose that water exclusion within the condensate microenvironment, coupled with protonation-driven RAC assembly, synergistically promotes dynamic arrest and aging. **Fig. 2i** illustrates a self-reinforcing cycle: initial arrest leads to droplet compaction, increasing hydrophobicity and glutamic acid protonation, which in turn drives further RAC assembly and solidification. This mechanistic insight into SynIDP aging provides a framework for tuning condensate morphology and optimizing peptide drug encapsulation.

## Controlling Condensate Aging to Sculpt Droplet and Filament Networks

Next, we aim to demonstrate that tunable solidification of SynIDP condensates enables precise control over gel morphology and feature size. To do so, we exploit two orthogonal and tunable variables— temperature and pH. The schema in **Fig. 3a** summarizes the temperature and pH pathway-dependent condensation and hierarchical assembly of the SynIDP. At pH 7.4, the SynIDP undergoes thermally driven LLPS into a liquid condensate (**Fig. 3b)**, and lowering the pH of this liquid condensate then drives its solidification into arrested condensate droplets with internal water-rich voids through microphase separation (**Fig. 3c)**. This internal microstructure has previously been observed in condensates following arrest [31, 32].

**Fig. 3:**
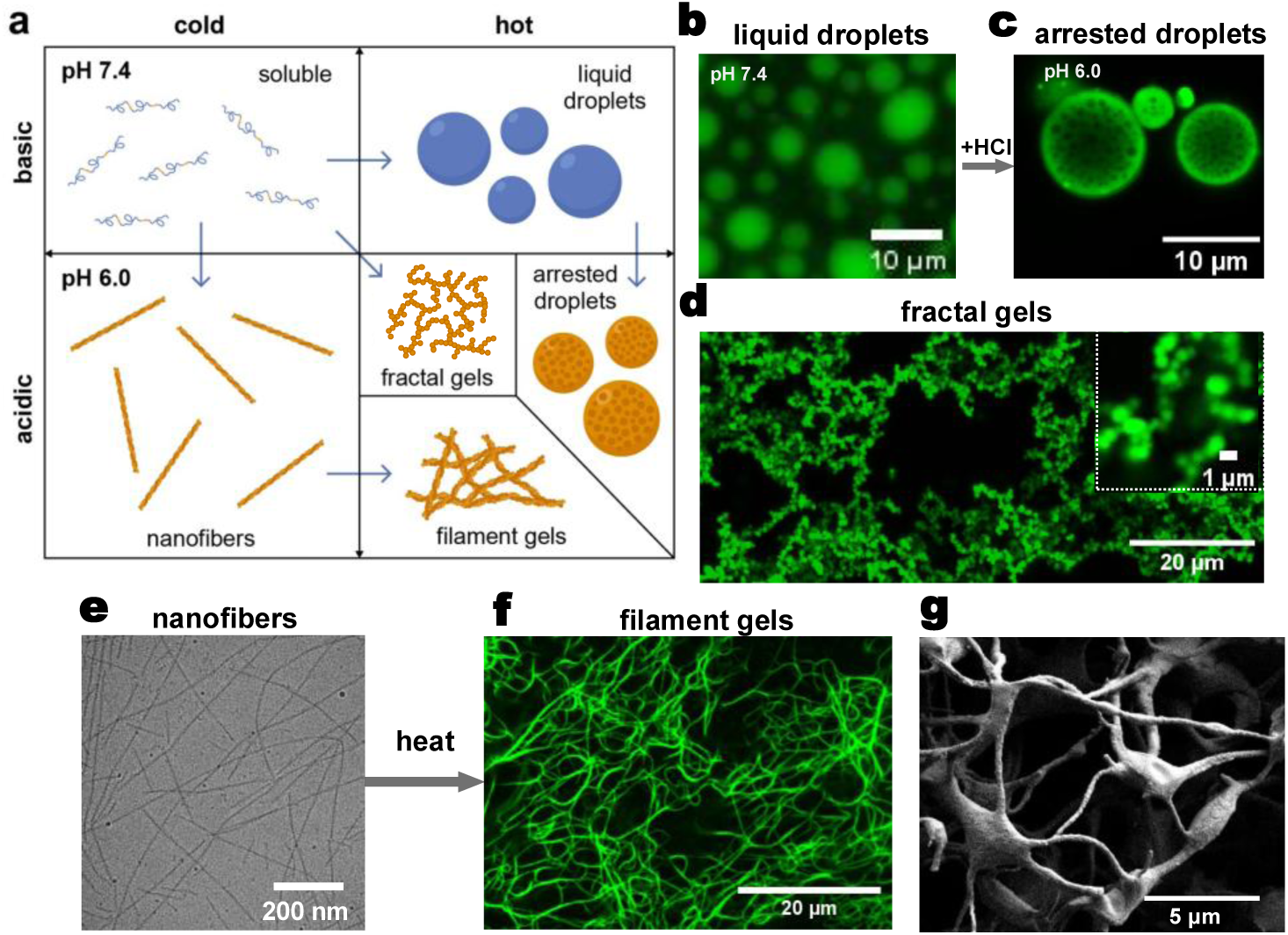
Distinct hierarchical assemblies from concerted transitions of synIDP. **a**, The temperature-pH phase diagram for pathway-dependent assembly to achieve each gel morphology from soluble synIDP; Heating and subsequent acidification produces arrested droplets, heating low pH samples leads to fractal gels with pH-dependent subunit sizes, and lowering pH to grow nanofibers before heating produces filament gels. **b,** Freshly prepared Alexafluor-488 labeled synIDP at pH 7.4 forms homogenous liquid droplets. **c,** Adjusting the pH to 6.0 with acid causes droplets to develop micro-phase separated structures. **d,** Heating fresh synIDP prepared at low pHs generates fractal gel networks: 2D projections of confocal z-stacks taken of synIDP transitioned at ph 5.2 with inserts showing aggregated microdroplet subunits. **e,** CryoTEM imaging of a pH 6.0 synIDP sample incubated for 3 days to grow high aspect ratio nanofibers. **f,** Z-stack projection of a filament gel formed by heating a pH 6 nanofiber sample after 3 days of incubation at 25 °C. **h,** SEM of a filament gel to show synIDP wetting of nanofiber bundles. Sample was prepared at pH 6 and has been flash frozen and desiccated.

The synIDP can also form fractal gels with defined subunit sizes by heating at acidic pHs below 6. Generally, liquid condensates undergo a nucleation and coalescence process in which small droplets grow over time. If solidification occurs concurrently while droplets are small and in the Brownian diffusion regime, they can stick together to form fractal structures via arrested coalescence. This process has been observed in other IDPs that show phase separation [33, 34] and recapitulated in silico by tuning gelation times [35, 36]. After the onset of temperature triggered condensation after heating to 37 °C in citrate-phosphate buffer at pH 6.0, “bead” (solid droplet) diameters of approximately 4 µm are observed due to their arrest occurring late in the coalescence process, on the time scale of 5-10 minutes (**Supplementary** Fig. 7). Lower pH accelerated arrest; at pH 5.2, bead diameters were <1 µm, indicating near-immediate solidification (**Fig. 3d)**. These observations align with the expected increase in the rate of RAC assembly as the pH approaches the solution pKa of glutamic acids, which should correspond with a faster solidification transition. Fractal networks composed of sub-micron to multiple micron-sized beads form when heating low pH samples, while heating neutral pH samples results in large droplets or mesoscale puddles (**Supplementary** Fig. 7). This approach of simultaneously heating to drive its temperature triggered phase separation and solidification as a function of pH provides a wide range of accessible droplet sizes.

If instead of heating, the pH of a solution of the SynIDP at room temperature is first reduced to pH 6 or lower, the SynIDP forms filaments, indicating pathway-controlled assembly. Incubation of pH 6 SynIDP in PBS at 30 °C — a temperature below the turbidity transition temperature of the SynIDP— overnight followed by heating to 45 °C, which is above its T_t_, produces micron-scale fiber bundles, as visualized by optical microscopy with ThT (**Supplementary** Fig. 8). Filaments are often composed of nanofiber bundles [37, 38], prompting us to investigate the assembly of nanofibers at low temperatures. Cryo-TEM reveals high aspect-ratio nanofibers that are ∼15 nm wide and hundreds of nanometers long (**Fig. 3e**).

Small angle X-ray scattering (SAXS) experiments were conducted to investigate the pH-dependent nanostructure of the system below its T_t_. As expected, the SAXS profile at neutral pH (**Supplementary** Fig. 9) indicates a random coil conformation characteristic of an unfolded protein, as confirmed by the Kratky plot’s open-ended shape [39]. A Guinier fit [40]yielded a radius of gyration (R_g_) of 7.5 nm. In contrast, the SAXS curve at pH 6 reveals fibrillar assembly, evidenced by a ∼−1 slope in the power-law region of the scattering curve [41]. Additionally, a Bragg peak at q = 0.05 Å⁻¹—absent at pH 7.4—corresponds to a real-space distance of 12.6 nm, which may reflect fibril spacing [42] (**Supplementary** Fig. 9). Complementary ThT fluorescence kinetics further support pH-dependent fibrillization: samples at pH 6 showed a rapid increase in fluorescence intensity, whereas those at pH ≥ 7 remained inert over an 18-h period (**Supplementary** Fig. 10).

Upon heating, nanofiber suspensions transitioned into filament gels composed of extended bundles (**Fig. 3f**). SEM imaging revealed smooth junctions, suggesting wetting by soluble protein polymer (**Fig. 3g**). Raising the pH immediately prior to heating induced rearrangement of the filament networks into micron-scale droplets that wet the scaffold (**Supplementary** Fig. 11). These assemblies were reversible with pH, as large bundles dissolved within minutes, while individual filaments required hours upon changing the pH to 7.4 (**Supplementary** Fig. 12). Together, these results reveal three distinct SynIDP morphologies—arrested droplets, fractal networks and filamentous bundles—that are accessible by manipulation of the temperature and pH of the SynIDP giving rise to pathway-controlled morphologies.

## Arrested SynIDP Droplets Survive in Simulated Gastric Acid

Based on the reversible transitions of the SynIDP, we devised the following strategy for drug encapsulation and enteric release (**Fig 4a**). First, a peptide drug is incubated with the synIDP to trigger supramolecular co-assembly. To form capsules of the drug that we hypothesize would survive in the pepsin-rich and low pH environment of the stomach, the mixture is heated to 45 °C to induce condensation into liquid condensate droplets. Next, dropping the pH of the droplet drives solidification of the condensate. We hypothesized that these arrested droplets would stay intact in the slightly acidic esophagus and highly acidic stomach, protecting the peptide drug cargo from proteolysis. Then, the coating would dissolve in the neutral to alkaline pH environment of the intestine to release the drug. We note that of the various morphologies available to us, we chose arrested droplets as the drug carrier, as its low surface area to volume ratio would minimize its contact with the protease-rich and acidic milieu of the stomach that rapidly degrades peptides and proteins.

**Fig. 4:**
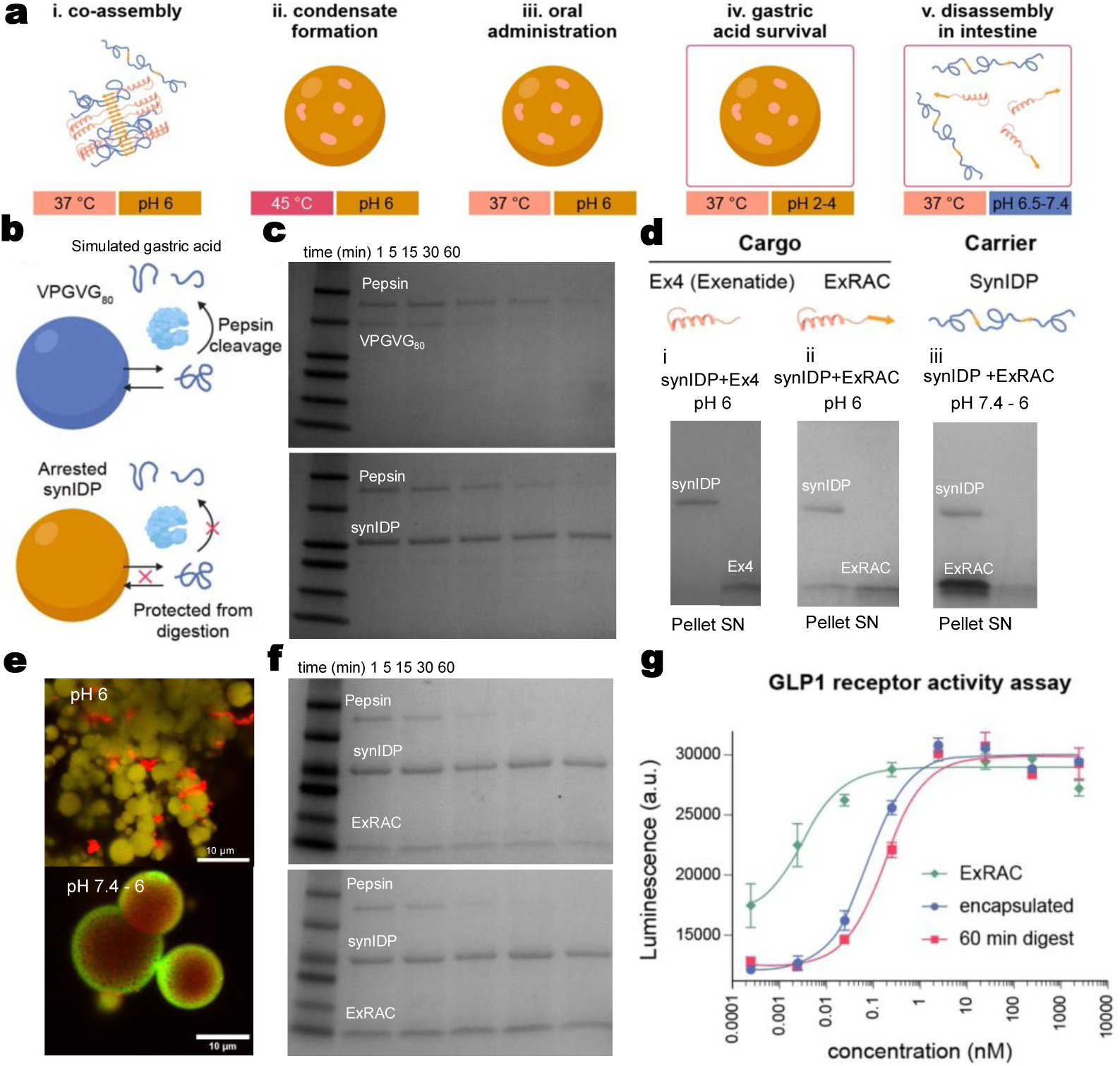
Encapsulation and protection of a peptide drug. **a**, The schema of assembly states of the synIDP and a peptide drug during preparation (**i-ii**), and in gastrointestinal environments (**iii-v**) for enteric delivery of cargo. Co-assembly allows for encapsulation in droplets upon heating, while low pH conditions keep the droplets stable below their equilibrium transition temperature during storage and upper GI transit. **b,** Schema of a liquid ELP droplet (top) and an arrested synIDP condensate (bottom) in simulated gastric acid (SGA). Liquid droplets are susceptible to digestion due to protein exchange with the dilute phase, while arrested condensates are more robust and mechanically stable. **c,** ELP condensates (top) and synIDP arrested condensates (bottom) were incubated in pH 1.5 SGA at 37 °C, from which aliquots were collected and analyzed by SDS-PAGE. **d,** The loading of peptide drug cargos into synIDP condensates was monitored by SDS-PAGE for synIDP transitioned with Ex4 at pH 6(**i**), synIDP transitioned with ExRAC at pH 6 (**ii**), and synIDP transitioned with ExRAC at pH 7.4 for 10 minutes, then adjusted to pH 6 **(iii**)**. e,** Overlayed confocal images of synIDP (green) and ExRAC (red) transitioned at pH 6 to yield fused droplets (top) or transitioned at pH 7.4 for 10 minutes, then adjusted to pH 6 (bottom), which creates void microstructures that contain ExRAC. **f,** SDS-PAGE gels corresponding to SGA digests of pH 6 (top) and pH 7.4 to 6 (bottom) formulations of synIDP with ExRAC. **g,** Quantification of GLP-1 receptor activation from incubating HEK cells with soluble ExRAC, ExRAC encapsulated then released from synIDP condensates, or encapsulated ExRAC that was digested for 60 minutes before release.

Before encapsulating a peptide drug in arrested condensates, we first investigated whether solidified SynIDP droplets would survive in simulated gastric conditions. To do so, we compared liquid ELP condensate droplets of a VPGVG_80_ sequence to an arrested SynIDP condensate after incubation in SGA (pH 1.5) with 1.2 mg/ml pepsin (**Fig. 4b,c**) as a function of incubation time. SDS-PAGE shows that liquid ELP condensate droplets could only withstand 15 min in SGA before the protein band disappears (**Fig. 4c, top**). For comparison, bovine serum albumin (BSA) and green fluorescent protein (GFP) were undetectable at the 1-minute time-point in SGA (**Supplementary** Fig. 13). In contrast, arrested SynIDP condensate droplets survive for at least 60 minutes in SGA (**Fig. 4c, bottom**), which corresponds to the transit time in the stomach for humans [43]. This confirms our hypothesis that a liquid condensate cannot survive in the stomach whereas an arrested solid condensate is resistant to the stomach environment.

Fluorescence-lifetime imaging microscopy (FLIM) offers a sensitive proxy for probing molecular interactions by detecting changes in fluorescence lifetime that reflect the local microenvironment [44, 45].We hypothesized that SynIDP condensates may exhibit increased microenvironmental density at lower pH, resulting in enhanced confinement of sequestered molecules. Consistent with this, we observed a pH-dependent decrease in fluorescence lifetime of Alexa-488–labeled SynIDP within the dense phase: from 3.458 ± 0.147 ns at pH 7.4 to 3.196 ± 0.130 ns at pH 6 **(Supplementary** Fig. 14**, top right**).

Notably, at pH 6, the fluorescence lifetime map revealed a core–shell architecture, with the condensate interface exhibiting a higher lifetime (∼3.4 ns) compared to the interior (∼2.9 ns) (**Supplementary** Fig. 14**, top left**). This spatial variation suggests that molecules within the condensate core are more confined than those near the periphery which may explain the ability of arrested condensates to dissolve from the interface in alkaline conditions. In contrast, condensates at pH 7.4 displayed a homogeneous fluorescence lifetime distribution (**Supplementary** Fig. 14).

In the dilute phase, Alexa-488–labeled SynIDP showed no significant pH-dependent variation, with lifetimes of 3.761 ± 0.336 ns at pH 7.4 and 3.822 ± 0.368 ns at pH 6 (**Supplementary** Fig. 14**, bottom**). These values align with previous reports indicating that Alexa-488 fluorescence lifetime is largely insensitive to pH [46]. Thus, the observed lifetime shift within condensates likely reflects increased non-radiative decay pathways associated with molecular confinement, rather than direct pH effects. This provides insight into the density and networking transitions in arrested droplets that may contribute to their resistance to degradation.

## Encapsulation of GLP1-RAs in Solidified Condensate Droplets

Next, we encapsulated peptide drugs within solidified SynIDP droplets. We selected exendin-4 (also known as exenatide), a 39-amino acid GLP-1 receptor agonist that stimulates insulin release, delays gastric emptying, and promotes satiety. To enhance co-assembly with SynIDP, we fused a copy of the CDC19 RAC domain to the flexible C-terminus of exendin-4, generating a modified peptide drug termed ExRAC. Both exendin-4 and ExRAC were cloned in *E. coli* as TEV-cleavable fusions to an elastin-like polypeptide (ELP) to facilitate expression and purification (**Supplementary** Fig. 15).

To test our hypothesis that the C-terminal RAC would promote co-assembly with the RAC domains embedded in SynIDP, we compared the encapsulation efficiency of ExRAC, exendin-4, and semaglutide. Peptide drugs were incubated with SynIDP at 37 °C for 2 hours at pH 6, then heated to 45 °C to trigger liquid–liquid phase separation (LLPS), followed by pelleting of the condensates by centrifugation. Analysis of supernatants and resuspended pellets revealed that exendin-4 was not encapsulated at pH 6 (**Fig. 4d, i**), and ExRAC showed some encapsulation under the same conditions (**Fig. 4d, ii**). When the mixture was adjusted to pH 7.4 immediately prior to heating, then acidified to pH 6 after 10 minutes of heating, we observed the highest encapsulation efficiency (**Fig. 4d, iii**).

Semaglutide was partially encapsulated at pH 6 (**Supplementary** Fig. 16), potentially due to hydrophobic interactions between its alkyl chain and the ELP domain, or due to its limited solubility at acidic pH, which may promote nucleation of condensates around semaglutide aggregates. In contrast, large droplets of SynIDP that were formed at pH 7.4 then acidified encapsulated up to 68% of ExRAC, as quantified by a bicinchoninic acid assay of fractionated droplets and supernatants.

Confocal imaging of SynIDP (green) and ExRAC (red) revealed distinct structural features depending on pH. At pH 7.4 acidified to pH 6, large condensates formed dense SynIDP coronas surrounding a peptide-rich core, with ExRAC penetrating the pores of dilute-phase SynIDP microstructures formed during arrest (**Fig. 4e, bottom**). At pH 6, droplet networks displayed co-localization in small droplets and peptide-rich aggregates (**Fig. 4e, top**), indicating differential encapsulation architectures.

After optimizing encapsulation conditions, we evaluated drug survival in SGA. Comparing pH 6 droplet networks to larger solidified condensates revealed that the latter protected most of the drug over 60 minutes, whereas smaller networks lost most of their payload during the same period (Fig. 4f). We attribute this to deeper burial of ExRAC within the condensate core and the lower surface area-to-volume ratio of larger droplets. Pepsin would need to erode more protein to access the drug in core–corona structures, whereas pH 6 networks expose a greater fraction of drug on the surface.

Semaglutide remained detectable in SGA for up to 30 minutes, as shown by SDS-PAGE (**Supplementary** Fig. 17**, left**). Because semaglutide does not migrate at its expected molecular weight on SDS-PAGE, we confirmed its integrity by MALDI-MS (**Supplementary** Fig. 17**, right**). Based on these findings, we selected large, solidified condensates for all subsequent experiments.

## Retention of Drug Bioactivity after Encapsulation and Simulated Gastric Digestion

We validated the bioactivity of ExRAC following encapsulation and simulated gastric digestion to confirm that the drug remains functional upon release from condensate droplets at slightly alkaline pH. Activity was assessed using a cellular assay in which mammalian cells engineered to express the GLP-1 receptor produce luciferase in response to receptor activation. After encapsulation in SynIDP and release via gentle mixing at pH 7.4, the EC₅₀ of ExRAC increased approximately 20-fold compared to the unencapsulated drug (**Fig. 4g**), likely reflecting partial drug loss during the encapsulation process. However, the reduction in activity after 60 minutes of simulated gastric digestion was only 60% relative to encapsulation without digestion, indicating that roughly 40% of the encapsulated drug survives gastric conditions and retains functional activity.

## Oral Drug Delivery in Mice

Building on our demonstration that a SynIDP condensate can encapsulate, protect, and release an active GLP-1 receptor agonist (GLP1-RA) under simulated gastric conditions, we conducted a preclinical study in mice with diet-induced obesity. We compared our top-performing encapsulation strategy against two widely used peptide drug delivery methods: oral administration with SNAC and subcutaneous injection. Oral formulations were dosed at 1000 nmol/kg—ten-fold higher than the subcutaneous dose—based on prior studies indicating a maximum oral bioavailability of approximately 10% for peptide drugs [47, 48].

For oral delivery, both the SynIDP-encapsulated and unencapsulated drugs were co-formulated with SNAC at pH 6 and administered daily via oral gavage. Throughout the study, mice had ad libitum access to a high-fat diet, mimicking real-world conditions in which oral therapeutics may be taken regardless of satiety. This represents a stringent test of efficacy, as oral semaglutide formulations with SNAC are typically administered on an empty stomach [49, 50].

We monitored weight loss over a 5-day treatment period and compared outcomes across four groups: daily *s.c.* ExRAC injection, oral delivery of the encapsulated drug, oral delivery of free drug, and vehicle control (**Fig. 5a, b**). By the study endpoint, both the oral encapsulated drug and *s.c.* injected drug groups exhibited significant weight loss (∼3% of starting body weight) relative to the vehicle control, as determined by two-way ANOVA with Tukey’s multiple comparisons performed daily. In contrast, the oral unencapsulated drug group showed highly variable weight loss, likely due to inconsistent stomach volume leading to differential drug degradation.

**Fig. 5:**
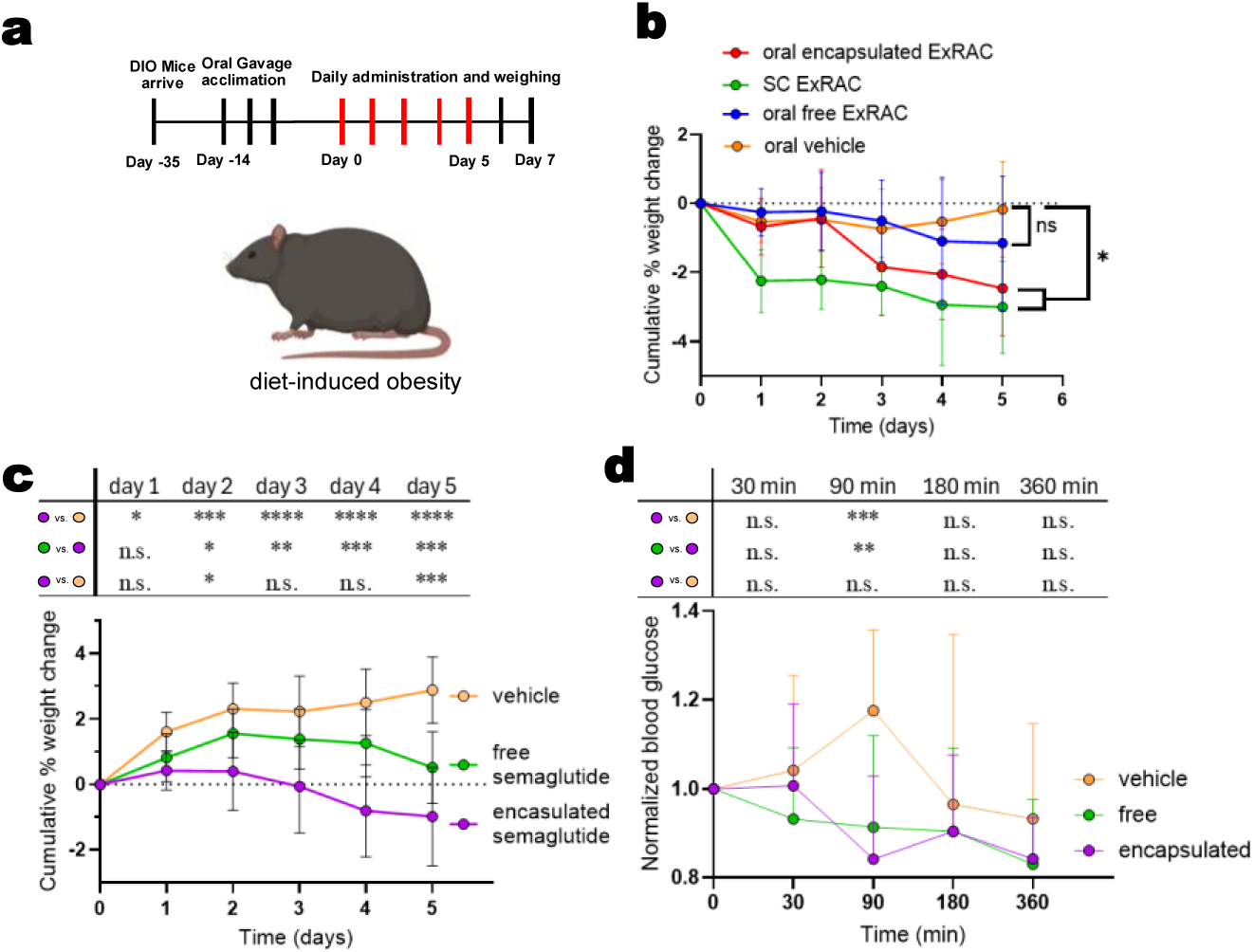
Efficacy of SynIDPs as enteric coatings in a mouse model of obesity. **a**, Obese mice were acclimated to oral gavage protocols and divided into groups with consistent average weights before being dosed with either oral or subcutaneous injections. **b,** Mouse weight change over 5 days of ExRAC administration show significant weight loss between both the oral SynIDP-encapsulated ExCDC and the oral vehicle control, and the injected ExCDC and the oral vehicle. **c,** Weight of mice administered with encapsulated semaglutide, free semaglutide, or vehicle were tracked over time; n=12 for therapeutic groups and n=8 for the the vehicle control. **d,** Blood glucose levels were measured daily for mice with orally administered semaglutide (encapsulated or free) or vehicle. Asterisks denote statistical significance (defined by * p<.05, ** p<.01, *** p<.001, **** p<.0001) as determined by 2-way ANOVA with Tukey’s test for with multiple comparison between groups at each timepoint.

To confirm the necessity of SNAC for intestinal absorption, we repeated the dosing regimen without SNAC. In all groups, this resulted in no measurable weight change over the same time period (**Supplementary** Fig. 18), reinforcing SNAC’s critical role in facilitating oral bioavailability.

Next, we encapsulated semaglutide in SynIDP and administered it orally to a larger cohort of mice using the same protocol. This yielded a markedly stronger distinction between the encapsulated, free, and vehicle control groups (**Fig. 5c**). We speculate that ExRAC undergoes rapid clearance from circulation, contributing to greater variability in outcomes across mice compared to a longer-circulating drug like semaglutide. The significantly greater weight loss observed in fed mice treated with SynIDP-encapsulated semaglutide, relative to free semaglutide, underscores the functional advantage of the SynIDP enteric coating.

To confirm that the enteric coating enhances drug bioavailability by protecting the payload during gastric transit, we investigated the pharmacokinetics of semaglutide and ExRAC in mice. Oral semaglutide formulations led to a significant reduction in blood glucose levels at 90 minutes post-administration compared to vehicle controls (**Fig. 5d**). Notably, encapsulated semaglutide exhibited a ∼1-hour delay in lowering glucose relative to free semaglutide, consistent with intestinal absorption following gastric passage rather than rapid uptake in the stomach. In contrast, blood glucose measurements in mice dosed with ExRAC showed no significant changes, except in the case of injected drug at 60 minutes post-delivery (Supplementary Fig. 19), likely due to ExRAC’s rapid systemic clearance and variability in gastric transit among fed animals.

We quantified circulating ExRAC concentrations in plasma using a competitive enzyme-linked immunosorbent assay (ELISA) against Exendin-4-binding antibodies. A standard curve was generated by spiking ExRAC into serum (**Supplementary** Fig. 20), which enabled us to determine blood concentrations across injected, encapsulated oral, and unencapsulated oral drug groups following administration (**Supplementary** Fig. 21). In mice treated with the encapsulated drug, ExRAC was largely undetectable at 1 hour but appeared at 2 hours, consistent with absorption in the lower intestine. Free ExRAC showed variable absorption: one mouse absorbed it in the stomach, two in the intestine, and one showed no detectable absorption. Injected ExRAC peaked at the earliest timepoint and gradually declined, with peak plasma concentrations for both injected and orally delivered drug consistently around 15 ng/µl. These results support the predicted ∼10% oral bioavailability and suggest that the 1-hour peak corresponds to ExRAC release in the neutral to alkaline environment of the intestine.

To investigate ExRAC disassembly from SynIDP droplets in the gastrointestinal tract, we imaged mouse GI tissues following administration of fluorescently labeled SynIDP (green) and ExRAC (red). At 30 minutes, the two components were predominantly co-localized in the stomach, with a small fraction entering the upper intestine (**Supplementary** Fig. 22). At later timepoints, additional drug and vehicle reached the upper intestine and cecum, where they began to spatially separate, indicating that drug release initiates in the upper intestine. Compared to free drug at 60 minutes, the encapsulated formulation traveled a shorter distance through the GI tract, potentially due to droplet adhesion. We also note that the lower stomach and duodenum contain elastase, which may contribute to condensate disassembly.

Overall, this proof-of-concept study in fed mice demonstrates that the SynIDP enteric coating can effectively protect and deliver peptide drugs through the gastrointestinal tract, supporting its potential as a versatile platform for oral delivery of biologics.

## Outlook

Enteric coatings have been used for nearly a century [51] to protect drugs from gastric acid and enable their release in the intestine. Modern synthetic copolymer enteric coatings employ two synergistic strategies for targeted intestinal delivery: phase separation to encapsulate drugs, and pH-sensitive chemistries that trigger dissolution [48]. These principles of stimulus-responsive encapsulation and release are mirrored in biomolecular condensates, making them compelling bioinspired templates for enteric coating design. Recent efforts have explored protein coacervates [52] and peptide analog droplets [53] for drug protection, though preclinical efficacy remains limited. Peptide coacervates can protect oligonucleotides from enzymatic degradation [54] and preserve protein activity in the dry state [55], further supporting their role in sequestration and protection. Importantly, a rich repertoire of latent, sequence-encoded functions can be harnessed to engineer modular multifunctional proteins for oral delivery, including supramolecular drug binding, specialized self-assembly cues, and control over enzymatic degradation.

Building upon these precedents, in this study, we design, characterize, and demonstrate the in vivo efficacy of the first engineered condensate-inspired protein polymer for enteric coating. While our system replicates the condensation, aging, and fibril-forming transitions observed in natural biomolecular condensates, it diverges morphologically from naturally-occurring IDPs. Specifically, we did not observe aging from spherical condensates into fibril bundles [22]; instead, fibril growth occurred in the dilute phase. Prior studies of FUS, hnRNPA1, and other condensate-forming proteins have reported fibril nucleation at interfaces and growth in the dilute phase [14, 56, 57], whereas our SynIDP condensates undergo dynamical arrest, as confirmed by FRAP, FLIM, and microrheology. Similar aging transitions have been described for engineered variants of the prion-like domain of hnRNPA1, where aging was driven by disorder-to-order transitions within the dense phase, leading to the formation of terminally viscoelastic Kelvin–Voigt solids [13].

Elucidating the molecular mechanisms underlying such distinct pathways of condensate aging remains an open question. In our case, this may stem from differences in the driving forces between native IDPs and SynIDPs. Our design principles—focused on encoding orthogonality and digestion resistance—intentionally omit several molecular interactions that drive native IDP phase separation, including charge complementarity, cation–π, and nonspecific π–π interactions. Condensates have also been reported to create electrochemical gradients that buffer their effective pH from their surroundings [58, 59]. This, combined with recent evidence that β-amyloid assembly is modulated by redox conditions in condensates [60], suggests there may be broader synergy between amyloid formation and the aging mechanism driven by microenvironment pKa shifts presented in this work.

Our enteric protein design leverages the modularity of its building blocks and a heuristic understanding of the factors governing phase separation in thermally responsive polypeptide fusions. The design process was non-trivial and iterative, with assembly behavior shown to be highly sensitive to sequence variations in the fiber-forming domains. The SynIDP enteric coating is expected to encapsulate and protect a range of therapeutics fused to cooperative self-assembly domains, such as the CDC19 RAC or hydrophobic alkyl chains. Moreover, we speculate that drug assemblies themselves could act as nucleation sites for condensation, enabling coatings to form around micelles, drug nanoparticles, or even cells. Ultimately, this condensate-based protein polymer may serve as a versatile platform not only for enteric delivery of diverse therapeutics but also for broader biotechnological applications in acidic physiological environments such as hypoxic wounds [64], lymph nodes [65], tumors [66], and the vaginal tract [67]. We anticipate that self-assembly could be tuned to occur upon heating from ambient to body temperature by modulating the hydrophobicity of the ELP domains, enabling the formation of injectable droplets or networked materials in response to local pH.

The selection of a modified exendin-4 peptide for this exploratory efficacy study was based on the hypothesis that co-assembly with SynIDP would enhance encapsulation and protection. However, exendin-4 lacks the extended circulation profiles of semaglutide and liraglutide, which are conjugated to alkyl chains. Exendin-4 has a reported half-life of 4–8 minutes [68], whereas semaglutide persists for days in humans [3, 69] and hours in mice[50]. This pharmacokinetic disparity may explain semaglutide’s superior performance relative to ExRAC, despite its lower encapsulation efficiency and gastric survival. In conclusion, while mice provide a suitable initial model for demonstrating preclinical efficacy of this novel protein-based enteric coating, further studies in additional species will be essential to evaluate its translational potential for oral drug delivery.

## Methods

### Cloning

Genes encoding the ELP fragments, CDC19 RAC, or peptide drugs were ordered from IDT (address) and placed into a custom pET24+ plasmid using Gibson Assembly, before concatenation to ELP genes by a process called Plasmid Reconstruction by Reversible Directional Ligation (PReRDL) [70]. Briefly, two plasmids with genes of interest were digested with Type IIs restriction endonucleases so that they contained complementary GG-CC overhangs at the intended site of fusion and were complementary at another point on each desired plasmid fragment. These fragments were purified by extraction of DNA from bands excised from agar gel electrophoresis, then ligated to assemble the gene. This process was repeated iteratively as needed until the full polypeptide gene was assembled. The sequence of the assembled genes are shown in **Supplementary Table S1**.

### Protein Expression and Purification

Proteins were expressed in BL21(DE3) *E. Coli* (New England Biolabs) transformed with pET-24+ bearing synthetic genes for the protein of interest and were purified by taking advantage of their reversible phase separation behavior, as described previously [18]. Cells were grown at 37 °C in 1 L of 2x YT medium in 4 L flasks and induced with Isopropyl β-D-1-thiogalactopyranoside overnight. Cells were spun down at 4,000 rcf for 15 min at 4 °C, and the cell pellet was resuspended in 20 mL of PBS, pH 8, before sonication on ice. Polyethyleneimine was added to the bacterial lysate to precipitate polynucleotides, following which the insoluble material in the lysate was separated by centrifugation at 15,000 rcf for 15 min at 4 °C. The supernatant was collected and heated to transition the protein polymers, which were then spun down in a heated centrifuge so that the pellet could be isolated. PBS (pH 8) was added to the pellet, which was then rotated gently overnight to resuspend. This constituted one round of inverse transition cycling. In total 3 rounds were performed until the protein was verified as pure by SDS-PAGE gel analysis and MALDI-TOF (**Supplementary** Fig. 23). Proteins were then dialyzed and lyophilized for storage.

Peptide drug fusions to ELPs were cleaved with Tobacco Etch Virus (TEV) protease (New England Biolabs) at the N-terminus of the peptide drug to leave two Alanine residues, which have been shown to increase Exendin circulation [71]. After overnight cleavage, the peptide drug was purified by one round of reverse ITC, followed by reverse immobilized metal affinity chromatography on a nickel column to remove free ELP, unreacted substrate, and TEV. The purified peptide drug was characterized by SDS-PAGE and MALDI-TOF. Finally, endotoxin was removed to a concentration below the FDA-recommended limit of less than 1 EU per maximum dose for oral or injected drugs [72] using Mustang E-membrane syringe-driven 0.2 µm filters (Pall Corporation) and verified a concentration, before dialysis and lyophilization.

### Brightfield and ThT Imaging

A 50x stock solution of Thioflavin T (Sigma-Aldrich) at a concentration of 2 mM ThT was prepared in PBS, pH 7.4, and filtered through a 2 µm syringe filter (Thermo Fisher) before being added to the protein samples at a final concentration of 0.2 µM. For imaging, precleaned glass slides were adorned with double-sided sticky tape lining the sample area to provide sufficient space for condensates to form between the slide and coverslip. A 3 mL droplet of sample was placed in the center of the slide without touching the tape to avoid wicking. Images were taken with a benchtop Axio Imager D2m microscope with a Linkam LTS120 heating stage at various temperatures. Brightfield, and green (excitation at 480 nm, emission at 520 nm, 2 s exposure), images were taken under 20x magnification and overlaid in ImageJ.

### Confocal Imaging and Fluorescence Recovery after Photobleaching (FRAP)

Confocal fluorescence imaging was performed on a Zeiss 710 inverted confocal microscope where the sample stage was heated to 40 °C with a custom incubator box (Zeiss). Samples of SynIDP were freshly resuspended at 0.4 mM in cold PBS or in 100 mM citrate phosphate buffer. For samples in PBS, (NH_4_)_2_SO_4_ was added to a final concentration of 80 mM to adjust to the T_t_ below 40 °C. The pH was adjusted to the desired value with HCl or NaOH, and in the case of the dynamic pH change experiments, we empirically determined how much acid or base to add to an equivalent volume of sample prior to imaging to toggle between the two pH conditions. Alexa Fluor 488-labeled protein was mixed with unlabeled protein for a final labeling percentage of 1%. Samples were added to a 384-well glass-bottom black Greiner plate and coated with mineral oil immediately prior to imaging. FRAP was performed following the methods and principles outlined in ref [73] Briefly, a small rectangular portion of a droplet or gel globule was photobleached using the Argon/2 488nm laser line at 100% intensity under 63x magnification, then the region of interest was imaged at 5 fps for 40 s to observe recovery. Spots were corrected for photofading using an unbleached region, then normalized pre-bleach intensity to 100% and post-bleach intensity to 0%. Three spots per sample were averaged to generate recovery curves.

### FRAP Curve Fitting

After plotting FRAP recovery curves in GraphPad Prism, GraphPad’s 1-phase nonlinear fitting function was used to generate exponential fitting equations for each curve. All curves with recovery >15% could be fit with R-squared accuracy values above 0.75, indicating a good fit [74]. The span of the fitted curves was used a proxy for maximum recovery, while the Tau values were compared to understand speed of recovery between samples.

### Temperature-Programmed UV-vis Absorbance Spectroscopy

Temperature-programmed UV-vis absorbance spectroscopy was performed on a UV-Vis spectrophotometer (Cary 3500, Agilent Technologies) with Peltier controlled thermal ramp capabilities of a 7 cuvette holder attachment. SynIDP samples were freshly resuspended at 4 °C at 10 mg/ml, then loaded into pre-cooled cuvettes. Temperature ramps were performed at 0.5 °C/min while monitoring the absorbance at 350 nm. To determine the amount of acid or base to add to samples to change the pH to the desired values, the samples were split into 2 aliquots with the same volume, and 0.5 N HCl or 0.5 N NaOH were titrated into one aliquot until the desired pH was reached. The same amount of NaOH or HCl was added into the sample in the cuvette of the spectrophotometer after reaching the final temperature of the heating cycle.

### Dynamic pH Switching after Heating-Induced Condensation

We added less than 1% v/ of acid (1N HCl) or base (1N NaOH) to SynIDP samples to adjust the pH of samples to the desired pH without diluting or physically disturbing the condensates. To determine the amount of acid or base to add, we split each sample into two identical aliquots. One was gradually adjusted to the desired pH while monitoring with ColorPhast pH 5-10 strips (ThermoFisher) so that the necessary volume could be calculated. Then, we added that volume to the sample undergoing monitoring for UV-Vis or FRAP during the experiment. Finally, we verified the measured aliquot post-testing.

### Sample Preparation for Video Particle Tracking Nanorheology and Fluorescence-Lifetime Imaging Microscopy (FLIM)

ELP and synIDP condensates were prepared by combining in the following order: peptide, water (Ambion), PBS (source), fluorescent 200 nm amine or carboxylate coated beads for VPT (Invitrogen), either Na_3_PO_4_ (Sigma-Aldrich) or HCl (source) for pH 7.4 or 6 respectively, and inducing condensates with (NH_4_)_2_SO_4_ (source) with the final concentrations being 10 mg/mL peptide, 1x PBS, 0.0002% beads (VPT only), and 125 mM (NH_4_)_2_SO_4_ unless otherwise specified. pH was determined by using a seven compact pH meter (Mettler Toledo) equipped with a micro pH probe (ThermoScientific) capable of measuring pH in as little as five microliters of sample. Concentrations of HCl and Na_3_PO_4_ were dependent upon the pH with ∼0.5 µL of 0.1 N HCl and ∼1 µL of 0.1 M Na_3_PO_4_ being sufficient for VPT.

Upon induction of condensates samples were sealed into a custom flow chamber made from 1”x3” slides, 18 x 18 mm #1.5 coverglass coated with sigmacote (Sigma-Aldrich) following the protocol provided by Sigma-Aldrich, and 3-5 layers of double-sided tape (3M). Five microliters of sample were applied to the cover glass before the chamber was sealed. Subsequently, mineral oil was added to the chamber to seal the sample and prevent evaporation.

### Video Particle Tracking Nanorheology

Imaging was conducted on a Zeiss Primovert inverted microscope with a 100x oil-immersion objective lens equipped with a temperature-controlled stage (Instec). Condensates were imaged immediately after preparation (∼5 minutes). For imaging, a Blackfly S USB3 CMOS camera

(Teledyne-FLIR) was used. The microscope was focused so that beads were in the interior of the condensate. Videos were collected for 1000 frames at a rate of 10 frames per second. Measurements were made for three independent sample preparations with a single field of view for aging experiments and at least three fields of view for salt dependence.

### Data Analysis for Video Particle Tracking Nanorheology

Videos of bead motion can be analyzed to get the mean square displacement (MSD) of the beads [23, 24]. The MSDs were analyzed to obtain the diffusion coefficients, nature of the diffusion, and the viscosity of the beads embedded inside the condensates. For tracking, we utilized the TrackMate plugin of ImageJ FIJI software [75] to extract two-dimensional trajectories of each bead within the condensates. To account for stage drift, we compute the center of mass vector *R* at each time point yielding the center of mass motion of the bead ensemble within the field of view. The calculation of the center of mass vector was done using particle velocities according to the following equation:

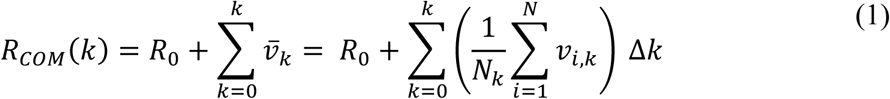

Here, *R*_0_ is the initial center of mass vector calculated by averaging the coordinates of all beads in the first frame 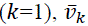 is the mean velocity of all particles in the frame *k*, *v*_*i*,*k*_ is the velocity of the particle *i* in frame *k* in units of µm/frame and *N*_*k*_ is the number of particles in the frame *k*. The quantity Δ*k* is the frame difference, which is 1 in the present case. To remove the stage drift, the center of mass is calculated and subtracted from the individual bead trajectories. This was used to calculate the ensemble-averaged mean-squared displacement (MSD) using:

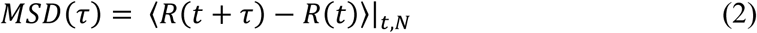

In Eq. (6), τ is the lag time. Next, we extracted the diffusion coefficient of the beads by fitting the MSD to the following equation:

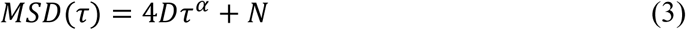

Here, *D* is the diffusion coefficient, α is the diffusivity exponent, and *N* is a term to account for the tracking noise. For fluids with terminally viscous behavior, α approaches 1 at long lag-times, which allows for the calculation of the terminal viscosity through the Stokes-Einstein equation:

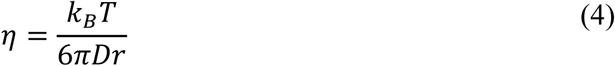

Here, *r* is the particle radius, *k*_*B*_is the Boltzmann coefficient, and *T* is the temperature in Kelvin. The averaged value was reported for the viscosity from the ELP condensates at different salt concentrations over three independently prepared samples. The error was estimated as the standard deviation.

### FLIM Data Collection and Analysis

Samples of ELP or SynIDP were prepared as described previously at 10 mg/ml protein in PBS with 125 mM (NH_4_)_2_SO_2_ at pH 7.4 or 6. Then fluorescence lifetime measurements were acquired using a laser-scanning confocal microscope designed for FLIM (ISS, Q2) equipped with a 488 nm laser and a PlanApo 63x water immersion objective (1.2 NA, Nikon) following a protocol described in a previous study [26, 73]. Briefly, to calibrate the FLIM module, rhodamine 110 was diluted to 1 mM and imaged following the manufacturer’s instructions. Samples were imaged shortly after induction (approximately 5 min), and three technical replicates were performed. Lifetimes were calculated using the Vistavision software (version 4.2, ISS), ensuring that Χ^2^ values of the fit were less than one. Maps of fluorescent lifetime in condensates were segmented by thresholding to separate the dilute and dense phases, and fitting was performed across the entire segmented image. Histograms of fluorescence lifetimes were generated using Vistavision, while GraphPad Prism 10 was used for generating the final fluorescence lifetime distribution plots along with the Gaussian fittings, which were used to provide the mean and standard deviation of the fluorescence lifetimes.

### Preparation of Nanofiber and Filament Samples

Lyophilized samples of SynIDP were freshly prepared in PBS (pH 7.4) or citrate-phosphate buffer (pH 6) and allowed to incubate at 25 °C in the same buffer or 37 °C for 24-72 h. Unlabeled SynIDP samples were prepared with ThT (0.2 µM) or mixed with 1% (w/w or v/v) Alexafluor-488 labeled SynIDP for imaging. For filament generation, samples were heated above the T_t_ in that buffer, and imaged by fluorescence/confocal fluorescence microscopy under 20x magnification on an Axio Imager D2m benchtop microscope, or with a Zeiss 720 inverted confocal microscope at 63x magnification.

### Cryo-TEM

SynIDP samples were resuspended in PBS, pH 6, at concentrations of 5mg/ml and imaged using TEM at cryogenic temperature. Samples were allowed to assemble at room temperature for 72 hours to generate nanofibers. Vitrified specimens were prepared on a copper grid coated with a perforated lacey carbon 300 mesh (Ted Pella Inc.). A typical 3 μL drop from the solution was applied to the grid in a controlled environment and blotted with a filter paper to form a thin liquid film of solution. The blotted samples were immediately plunged into liquid ethane at its freezing point (−183 °C). The procedure was performed automatically in the Plunger (Leica EM GP). The vitrified specimens were then transferred into liquid nitrogen for storage. The samples were studied using a FEI Talos F200C TEM, 200 kV maintained at liquid nitrogen temperature; and images were recorded by FEI Ceta16M camera (4k × 4k CMOS sensor) at low dose conditions, to minimize electron beam radiation damage

### Thioflavin T assay

Lyophilized SynIDP was dissolved in PBS at 1 mg/ml and mixed with Thioflavin T (ThT) (Thermo Fisher) at a final concentration of 40 nM. One SynIDP sample was prepared at pH 5.8 and another at pH 7.4, and these samples were mixed to generate intermediate pH samples for testing. Samples were added to black clear bottom 96 well plates (Greiner) and the fluorescence (Ex = 450 nm, Em = 480 nm) was measured using a Molecular Devices SpectraMax iD5 plate reader. The plates were incubated at 37 °C with intermittent orbital mixing and measured every 5 min for 18 h.

### SAXS

SAXS data were collected at the 16ID-LiX Beamline (National Synchrotron Light Source II, Brookhaven National Laboratory, Upton, NY). The X-ray wavelength was 0.8189 Å, and two setups (small– and wide-angle X-ray scattering) were used simultaneously to cover scattering q ranges of 0.006 ≤ q ≤ 3.19 Å−1, where q = (4π/λ)sinθ, with 2θ representing the scattering angle and λ indicating the X-ray wavelength.

SynIDP samples were measured at temperature of 25°C using custom-made flat fixed cells with two mica windows. For each measurement using a fixed cell, both empty cells and matching buffer were measured before solution scattering measurements for proper background subtraction. Twenty repeated measurements were taken on various parts of the sample with a 0.5s exposure time to avoid radiation damage. Radial averaging and q-conversion of data were performed using standard software to merge data from the two detectors used in SAXS and WAXS measurements. Transmission correction and background subtraction were performed to minimize the intensity of the hydrogen bond from water. The final scattering profiles were obtained by averaging the repeated measurements and subtracting the empty cell and blank solvent contributions. Data processing was performed using LiXTools (https://github.com/NSLS-II-LIX/lixtools) and the Python package py4xs (https://github.com/NSLS-II-LIX/py4xs) in Jupyter Notebook. Final scattering profiles were also manually validated using the Irena package for analysis of small angle scattering data. [76]

### Scanning Electron Microscopy

100 µl of SynIDP nanofiber suspensions at pH 6 were prepared according to previous descriptions and added to a custom mold for SEM sample processing. The suspension was heated for 15 minutes to trigger filamentous gel assembly, then the mold was flash frozen in liquid nitrogen to preserve its features. Following freezing, the sample was lyophilized, then carefully removed from the mold. The lyophilized sample was gold-sputtered with a Denton Desk V sputter coater. Electron micrographs were recorded with a Hitachi TM3030Plus Tabletop SEM.

### Drug Encapsulation

ExRAC and semaglutide (TargetMol Chemicals) were suspended at 5 mg/ml in 100 mM citrate phosphate buffer at a pH between 5.2 and 7.4 and mixed 1:1 (v/v) with freshly prepared 10 mg/ml SynIDP in deionized water at room temperature for at least 2 h to form nanofibers. The mixture was then placed on a heat block at 42 °C for 10 min to trigger the phase transition of the SynIDP. If the buffer was not already at pH 6 or lower, it was adjusted to pH 6 to arrest the condensate transition. The mixture was then centrifuged at 3,000 rcf for 4 min at 40 °C and the pellet was resuspended in an equivalent volume of pH 6 PBS for imaging on microscope slides, for SGA experiments, or for preparation for oral administration to mice.

### Simulated Gastric Acid

Simulated gastric acid (SGA) was prepared according to previous descriptions [77]. Immediately before digestion, 1.2 mg/ml of lyophilized porcine pepsin was dissolved in SGA buffer, which consisted of 35 mM and 0.1 N HCl adjusted the pH to 1.5 at 37 °C. Protein samples were added to this SGA mixture in a 1:7 ratio, then 15 µl aliquots were taken at 1, 5, 15, 30, and 60 min. These aliquots were quenched with 200 mM sodium bicarbonate buffer (pH 9) then heat inactivated at 80 °C for at least 10 min. Samples were analyzed by SDS-PAGE for survival of the protein condensates and their cargos.

### Quantification of GLP-1R Activation by cAMP-Responsive Luciferase Assay

To determine the activity of ExRAC in the digested or undigested fusion protein, a HEK293 cell line stably expressing the GLP-1 receptor and a cAMP-inducible luciferase reporter gene was used. Cells were plated at 1 × 10⁵ cells/cm² in 96-well plates and incubated overnight. In parallel, ExRAC samples were pretreated with dipeptidyl peptidase-4 (DPP4, ProSpec-Tany) at a 1:500 molar ratio (DPP4:ExRAC) and incubated overnight at room temperature to cleave the leader peptide and reveal the biologically active N-terminus. If the ExRAC sample was coated with SynIDP, the mixture was adjusted to pH 7.4 with mild base and gently mixed until the sample was optically clear. The following morning, cells were treated with serial dilutions of GLP-1R agonist and incubated for 5 h. After incubation, media were replaced with Bright-Glo luciferase substrate (Promega, Madison, WI), and the luminescence was measured using a Victor X3 plate reader (Perkin Elmer). Signals were normalized against vehicle-treated controls, and dose-response curves were fit using GraphPad Prism 8 to calculate EC₅₀ values via a three-parameter logistic fit.

### Mouse Studies

Mouse experiments were approved by the Duke University Institutional Animal Care and Use Committee (IACUC) and were conducted in accordance with protocol A059-24-03. Male 9-week-old C57BL/6J mice (the Jackson Laboratory) with diet-induced obesity (DIO) were kept on a 60% fat diet (Bio-serv) for the course of the study. They were acclimated to handling and oral gavage with 18 gauge, 1.5-inch animal feeding needles prior to the start of treatment. They were separated into groups with an approximately average weight of 40 g and kept in mixed cages. During the study, fresh peptide drug (semaglutide or ExRAC) and SynIDP were prepared in sterile and endotoxin-free buffer and encapsulated for oral dosing each day following the methods for encapsulation described above. After encapsulation, protein droplet suspensions were mixed with SNAC that was adjusted to pH 6 for a per-mouse dose of 30mg/kg. Mice in the subcutaneous group received a daily 0.1 ml injection of free ExRAC at 100nm/kg in PBS (n=5). Oral dosing groups received encapsulated ExRAC (n=7) or semaglutide (n=12) at 1000 nM/kg, unencapsulated ExRAC (n=7) or semaglutide (n=12), or a vehicle control of SynIDP and SNAC (n=5 for ExRAC study, n=8 for seamgltuide study). Mice were weighed daily, then dosed at 2 pm for 5 consecutive days to monitor weight loss. Their weights were normalized to the start of the study, and results were plotted in GraphPad Prism and analyzed by 2-way ANOVA with multiple comparison for significance by Tukey’s multiple comparison test for mixed factor comparison between groups at each timepoint. Asterisks denote significance as defined by by * p<.05, ** p<.01, *** p<.001, **** p<.0001.

### Blood Glucose Measurement

On the day of blood glucose measurement, mice had their high fat diet removed 2 hours prior to dosing to lower the variability of glucose levels. Mice were dosed as described above using oral gavage of encapsulated drug, free drug, or vehicle, all formulated with SNAC at pH 6 in sterile PBS. After dosing, high fat diet was immediately provided to the mice. Blood was sampled using a lancet at the tail vein. The first droplet was washed away, then 0.3 µl of serum was absorbed onto the test strip. Blood glucose was measured with an AlphaTrak3 monitor and strips, which was calibrated using the provided reference solution before each timepoint. For analysis, the initial blood glucose measurement was used as the baseline for each mouse to normalize blood glucose foreach subsequent time point. Results were plotted and analyzed in GraphPad Prism and analyzed by 2-way ANOVA by Tukey’s multiple comparison test for significance (defined as * p<.05, ** p<.01, *** p<.001) between groups at each timepoint.

### IVIS Imaging

The high fat diet was replaced with a low fluorescence diet (Bio-serv Rodent Diet AIN-93M) two days prior to imaging. Mice were dosed with SynIDPencapsulated or unencapsulated drug as described previously, in which ExRAC was N-terminally labeled with Alexafluor-647 and SynIDP, if present, was N-terminally labeled with Alexafluor-488. After dosing, mice were sacrificed and their entire gastrointestinal tracts were dissected at set time points for imaging. Imaging was performed on an IVIS Lumina III system with green fluorescence images taken with 460 nm excitation and 520 nm emission filter,s and red images taken with 640 nm excitation and 670 nm emission filters, which were then analyzed and quantified using LivingImage software and ImageJ.

### ELISA

After dosing, 10 µl of blood was collected from the tail vein at multiple time points. Whole blood was centrifuged at 2000 rpm for 5 min and serum was stored at –80 °C until ELISA analysis. Serum concentrations were determined using an Exendin-4 (Heloderma suspectum) competition ELISA kits purchased from Creative Diagnostics, in which a standard curve using ExRAC doped into serum was generated for comparison to the Exendin-4 calibration samples. Each sample was diluted 100x from serum and then incubated with primary antibody and biotinylated peptide, incubated with streptavidin horseradish peroxidase, washed with assay buffer, reacted with tetramethylbenzidine substrate, and then quantified according to the kit instructions. Absorbance intensities 450 nm were measured on a Molecular Devices SpectraMax iD5 plate reader and fit to the standard curve to determine concentrations. Results were plotted in GraphPad Prism.

### MALDI-TOFMS

1 µL of sample (100uM protein in 150 mM PBS was mixed with 3 µL of a saturated solution of Sinapinic acid (SA) matrix and deposited on a ground steel MALDI plate. Samples were analyzed on a Bruker Autoflex LRF matrix-assisted laser desorption ionization tandem time of flight mass spectrometer (MALDI-TOF MS).

### Author Contributions

M.N. and A.C. conceived and planned the research. M.N., P.S., Y.S., A.S., G.W. X.L. J.Z., E.P.,

L.F., T.S.M., S.D., N.T., J.S., and J.M. performed the experiments. M.N., J.Z., J.S., and J.M. produced and purified the proteins. M.N., Y.S., P.S., L.F., E.P., and S.D. performed experiments to characterize the assembly states and MALDI-TOF verification of molecular weight. M.N. and N.T. performed SGA testing. P.S. performed cell culture and GLP1-RA assays. M.N. and X.L. carried out the mouse studies. G.W., A.S., and T.S.M. performed nanorheology and FLIM studies and interpreted results. M.N. and A.C. interpreted the results and wrote the manuscript. All authors provided feedback, discussion, and edits to the manuscript. P.R.B and A.C. were responsible for project administration, supervision, and funding acquisition.

## Acknowledgements

The HEK cells with GLP1 receptor activity were generously provided by Timothy Kieffer (University of British Columbia). We are grateful to the Duke Shared Materials Instrumentation Facility (SMIF) staff and resources for training and access. We thank Ben Carlson for FRAP assistance, Mark Walters for SEM and confocal assistance, and Talmage Tyler for sputter coating assistance. We thank Einat Nativ and the Ilse Katz Institute for Nano-Science and Technology at Ben Gurion University for CryoTEM samples preparation and imaging. Joel Collier and Emily Roe consulted on oral gavage and GI imaging study design. David D’Alessio consulted on mouse study design and data interpretation.

The SAXS data were collected at LiX beamline, which is part of the Center for BioMolecular Structure (CBMS). The beamline is primarily supported by the NIH, National Institute of General Medical Sciences (NIGMS), through a P30 Grant (P30GM133893), and by the DOE Office of Biological and Environmental Research (KP1605010). LiX also received additional support from NIH Grant S10 OD012331. As part of NSLS-II, a national user facility at Brookhaven National Laboratory, work performed at the CBMS is supported in part by the U.S. Department of Energy, Office of Science, Office of Basic Energy Sciences Program under contract number DE-SC0012704. This work was supported by the Intramural Research Program of the NIH, the National Cancer Institute (NCI) (ZIA BC 010379, ZIC BC 011535, and ZIA BC 011669 to Y-X.W.). The work performed in the Banerjee laboratory was supported by National Institutes of Health grant R35 GM138186 (to PRB) and St. Jude Children’s Research Collaborative on the Biology and Biophysics of RNP Granules (to PRB). The work performed in the Chilkoti lab was supported by the Air Force Office of Scientific Research MURI program grant (FA9550-20-1-0241) and a ‘Beyond the Horizons’ award from the Pratt School of Engineering at Duke University. The funders had no role in study design, data collection and analysis, decision to publish or preparation of the manuscript.

## References

1. Xiao, W., et al., Advance in peptide-based drug development: delivery platforms, therapeutics and vaccines. Signal Transduct Target Ther, 2025. 10(1): p. 74.

2. Yap, M.K.K. and N. Misuan, Exendin-4 from Heloderma suspectum venom: From discovery to its latest application as type II diabetes combatant. Basic Clin Pharmacol Toxicol, 2019. 124(5): p. 513–527.

3. Granhall, C., et al., Pharmacokinetics, Safety and Tolerability of Oral Semaglutide in Subjects with Renal Impairment. Clin Pharmacokinet, 2018. 57(12): p. 1571–1580.

4. Muttenthaler, M., et al., Trends in peptide drug discovery. Nat Rev Drug Discov, 2021. 20(4): p. 309–325.

5. Wang, J., et al., A Molecular Grammar Governing the Driving Forces for Phase Separation of Prion-like RNA Binding Proteins. Cell, 2018. 174(3): p. 688–699 e16.

6. Alberti, S. and A.A. Hyman, Biomolecular condensates at the nexus of cellular stress, protein aggregation disease and ageing. Nat Rev Mol Cell Biol, 2021. 22(3): p. 196–213.

7. Luo, F., et al., Atomic structures of FUS LC domain segments reveal bases for reversible amyloid fibril formation. Nat Struct Mol Biol, 2018. 25(4): p. 341–346.

8. Gui, X., et al., Structural basis for reversible amyloids of hnRNPA1 elucidates their role in stress granule assembly. Nat Commun, 2019. 10(1): p. 2006.

9. Guenther, E.L., et al., Atomic structures of TDP-43 LCD segments and insights into reversible or pathogenic aggregation. Nat Struct Mol Biol, 2018. 25(6): p. 463–471.

10. Frey, L., et al., A structural rationale for reversible vs irreversible amyloid fibril formation from a single protein. Nat Commun, 2024. 15(1): p. 8448.

11. Knowles, T.P. and R. Mezzenga, Amyloid Fibrils as Building Blocks for Natural and Artificial Functional Materials. Adv Mater, 2016. 28(31): p. 6546–61.

12. Linsenmeier, M., et al., Dynamic arrest and aging of biomolecular condensates are modulated by low-complexity domains, RNA and biochemical activity. Nat Commun, 2022. 13(1): p. 3030.

13. Alshareedah, I., et al., Sequence-specific interactions determine viscoelasticity and aging dynamics of protein condensates. Nat Phys, 2024. 20(9): p. 1482–1491.

14. Das, T., et al., Tunable metastability of condensates reconciles their dual roles in amyloid fibril formation. Mol Cell, 2025. 85(11): p. 2230–2245 e7.

15. McDaniel, J.R., D.C. Radford, and A. Chilkoti, A unified model for de novo design of elastin-like polypeptides with tunable inverse transition temperatures. Biomacromolecules, 2013. 14(8): p. 2866–72.

16. Meyer, D.E. and A. Chilkoti, Quantification of the effects of chain length and concentration on the thermal behavior of elastin-like polypeptides. Biomacromolecules, 2004. 5(3): p. 846–51.

17. Cho, Y., et al., Effects of Hofmeister anions on the phase transition temperature of elastin-like polypeptides. J Phys Chem B, 2008. 112(44): p. 13765–71.

18. MacEwan, S.R., W. Hassouneh, and A. Chilkoti, Non-chromatographic purification of recombinant elastin-like polypeptides and their fusions with peptides and proteins from Escherichia coli. J Vis Exp, 2014(88).

19. Vrhovski, B. and A.S. Weiss, Biochemistry of tropoelastin. Eur J Biochem, 1998. 258(1): p. 1–18.

20. Dunn, B.M., Structure and mechanism of the pepsin-like family of aspartic peptidases. Chem Rev, 2002. 102(12): p. 4431–58.

21. Cereghetti, G., et al., An evolutionarily conserved mechanism controls reversible amyloids of pyruvate kinase via pH-sensing regions. Dev Cell, 2024. 59(14): p. 1876–1891 e7.

22. Patel, A., et al., A Liquid-to-Solid Phase Transition of the ALS Protein FUS Accelerated by Disease Mutation. Cell, 2015. 162(5): p. 1066–1077.

23. Alshareedah, I., et al., Programmable viscoelasticity in protein-RNA condensates with disordered sticker-spacer polypeptides. Nature communications, 2021. 12(1): p. 6620.

24. Alshareedah, I., et al., Determinants of viscoelasticity and flow activation energy in biomolecular condensates. Science Advances, 2024. 10(7): p. eadi6539.

25. Mahendran, T.S., et al., Homotypic RNA clustering accompanies a liquid-to-solid transition inside the core of multi-component biomolecular condensates. Nature Chemistry, 2025: p. 1–11.

26. Mahendran, T.S., et al., Decoupling Phase Separation and Fibrillization Preserves Activity of Biomolecular Condensates. bioRxiv, 2025.

27. Alshareedah, I., G.M. Thurston, and P.R. Banerjee, Quantifying viscosity and surface tension of multicomponent protein-nucleic acid condensates. Biophysical journal, 2021. 120(7): p. 1161–1169.

28. Wake, N., et al., Expanding the molecular grammar of polar residues and arginine in FUS phase separation. Nat Chem Biol, 2025. 21(7): p. 1076–1088.

29. Mehler, E.L., et al., The role of hydrophobic microenvironments in modulating pKa shifts in proteins. Proteins, 2002. 48(2): p. 283–92.

30. Urry, D.W., S. Peng, and T. Parker, Delineation of electrostatic– and hydrophobic-induced pKa shifts in polypentapeptides: the glutamic acid residue. Journal of the American Chemical Society, 2002. 115(16): p. 7509–7510.

31. Erkamp, N.A., et al., Spatially non-uniform condensates emerge from dynamically arrested phase separation. Nat Commun, 2023. 14(1): p. 684.

32. Milligan, J.J., et al., Controlling Release Kinetics of an Adjuvant from a Depot Improves the Efficacy of Local Immunotherapy in Metastatic Cancer. Adv Sci (Weinh), 2025: p. e03591.

33. Lee, D.S.W., et al., Size distributions of intracellular condensates reflect competition between coalescence and nucleation. Nat Phys, 2023. 19(4): p. 586–596.

34. Firoozmand, H., B.S. Murray, and E. Dickinson, Fractal-type particle gel formed from gelatin + starch solution. Langmuir, 2007. 23(8): p. 4646–50.

35. Garaizar, A., et al., Kinetic interplay between droplet maturation and coalescence modulates shape of aged protein condensates. Sci Rep, 2022. 12(1): p. 4390.

36. Garaizar, A., et al., Aging can transform single-component protein condensates into multiphase architectures. Proc Natl Acad Sci U S A, 2022. 119(26): p. e2119800119.

37. Kanno, T., et al., Association of thin filaments into thick filaments revealing the structural hierarchy of amyloid fibrils. J Struct Biol, 2005. 149(2): p. 213–8.

38. Prince, E., et al., Nanocolloidal hydrogel mimics the structure and nonlinear mechanical properties of biological fibrous networks. Proc Natl Acad Sci U S A, 2023. 120(51): p. e2220755120.

39. Receveur-Bréchot, V. and D. Durand, How random are intrinsically disordered proteins? A small angle scattering perspective. Current Protein and Peptide Science, 2012. 13(1): p. 55–75.

40. Guinier, A., et al., Small-angle Scattering of X-rays. 1955: Wiley New York.

41. Glatter, O., Small angle X-ray scattering. (No Title), 1982.

42. Amiya, T. and T. Tanaka, Phase transitions in crosslinked gels of natural polymers. Macromolecules, 1987. 20(5): p. 1162–1164.

43. Maurer, J.M., et al., Gastrointestinal pH and Transit Time Profiling in Healthy Volunteers Using the IntelliCap System Confirms Ileo-Colonic Release of ColoPulse Tablets. PLoS One, 2015. 10(7): p. e0129076.

44. Datta, R., et al., Fluorescence lifetime imaging microscopy: fundamentals and advances in instrumentation, analysis, and applications. J Biomed Opt, 2020. 25(7): p. 1–43.

45. Ma, J., et al., Design and Application of Fluorescent Probes to Detect Cellular Physical Microenvironments. Chemical Reviews, 2024. 124(4): p. 1738–1861.

46. Scalia, G. and F. Scheffold, Lifetime of fluorescent dye molecules in dense aqueous suspensions of polystyrene nanoparticles. Optics Express, 2015. 23(23): p. 29342–29352.

47. Twarog, C., et al., Intestinal Permeation Enhancers for Oral Delivery of Macromolecules: A Comparison between Salcaprozate Sodium (SNAC) and Sodium Caprate (C(10)). Pharmaceutics, 2019. 11(2).

48. Liu, L., et al., pH-Responsive carriers for oral drug delivery: challenges and opportunities of current platforms. Drug Deliv, 2017. 24(1): p. 569–581.

49. Aroda, V.R., et al., Efficacy and safety of once-daily oral semaglutide 25 mg and 50 mg compared with 14 mg in adults with type 2 diabetes (PIONEER PLUS): a multicentre, randomised, phase 3b trial. Lancet, 2023. 402(10403): p. 693–704.

50. Rakhat, Y., et al., Oral Semaglutide under Human Protocols and Doses Regulates Food Intake, Body Weight, and Glycemia in Diet-Induced Obese Mice. Nutrients, 2023. 15(17).

51. Brown, T.D., K.A. Whitehead, and S. Mitragotri, Materials for oral delivery of proteins and peptides. Nature Reviews Materials, 2019. 5(2): p. 127–148.

52. Chen, S., et al., Extreme pH Tolerance in Peptide Coacervates Mediated by Multivalent Hydrogen Bonds for Enzyme-Triggered Oral Drug Delivery. J Am Chem Soc, 2025. 147(11): p. 9704–9715.

53. Chen, K., et al., Biomolecular Condensates Based on Amino Acid for Enhancing Oral Bioavailability and Therapeutic Efficacy of Hydrophobic Drugs. ACS Appl Mater Interfaces, 2024. 16(43): p. 58370–58378.

54. Rose Galvan, A., et al., Peptide Coacervates Can Protect Sequestered Oligonucleotides from Nucleases and Release Them for Transcription and Translation. Biomacromolecules, 2025.

55. Dave, D.R., et al., Adaptive peptide dispersions enable drying-induced biomolecule encapsulation. Nat Mater, 2025. 24(9): p. 1465–1475.

56. Choi, C.H., et al., Condensate interfaces can accelerate protein aggregation. Biophys J, 2024. 123(11): p. 1404–1413.

57. Shen, Y., et al., The liquid-to-solid transition of FUS is promoted by the condensate surface. Proc Natl Acad Sci U S A, 2023. 120(33): p. e2301366120.

58. Yu, W., et al., Aging-dependent evolving electrochemical potentials of biomolecular condensates regulate their physicochemical activities. Nat Chem, 2025. 17(5): p. 756–766.

59. Dai, Y., et al., Interface of biomolecular condensates modulates redox reactions. Chem, 2023. 9(6): p. 1594–1609.

60. Chen, M.W., et al., Transition-State-Dependent Spontaneous Generation of Reactive Oxygen Species by Abeta Assemblies Encodes a Self-Regulated Positive Feedback Loop for Aggregate Formation. J Am Chem Soc, 2025. 147(10): p. 8267–8279.

61. Malay, A.D., et al., Spider silk self-assembly via modular liquid-liquid phase separation and nanofibrillation. Science Advances, 2020. 6(45): p. eabb6030.

62. Mozhdehi, D., et al., Genetically encoded lipid-polypeptide hybrid biomaterials that exhibit temperature-triggered hierarchical self-assembly. Nat Chem, 2018. 10(5): p. 496–505.

63. Roberts, S., et al., Injectable tissue integrating networks from recombinant polypeptides with tunable order. Nat Mater, 2018. 17(12): p. 1154–1163.

64. Sim, P., et al., Influence of Acidic pH on Wound Healing In Vivo: A Novel Perspective for Wound Treatment. Int J Mol Sci, 2022. 23(21).

65. Wu, H., et al., T-cells produce acidic niches in lymph nodes to suppress their own effector functions. Nat Commun, 2020. 11(1): p. 4113.

66. Feng, Q., et al., Severely polarized extracellular acidity around tumour cells. Nat Biomed Eng, 2024. 8(6): p. 787–799.

67. O’Hanlon, D.E., R.A. Come, and T.R. Moench, Vaginal pH measured in vivo: lactobacilli determine pH and lactic acid concentration. BMC Microbiol, 2019. 19(1): p. 13.

68. Parkes, D., et al., Pharmacokinetic actions of exendin-4 in the rat: Comparison with glucagon-like peptide-1. Drug Development Research, 2001. 53(4): p. 260–267.

69. Hall, S., D. Isaacs, and J.N. Clements, Pharmacokinetics and Clinical Implications of Semaglutide: A New Glucagon-Like Peptide (GLP)-1 Receptor Agonist. Clin Pharmacokinet, 2018. 57(12): p. 1529–1538.

70. McDaniel, J.R., et al., Recursive Directional Ligation by Plasmid Reconstruction Allows Rapid and Seamless Cloning of Oligomeric Genes. Biomacromolecules, 2010. 11(4): p. 944–952.

71. Yao, J.F., et al., Metabolism of Peptide Drugs and Strategies to Improve their Metabolic Stability. Curr Drug Metab, 2018. 19(11): p. 892–901.

72. Administration, U.S.F.a.D., Guidance for Industry: Pyrogen and Endotoxins Testing: Questions and Answers.

73. Alshareedah, I., T. Kaur, and P.R. Banerjee, Methods for characterizing the material properties of biomolecular condensates. Methods Enzymol, 2021. 646: p. 143–183.

74. Taylor, N.O., et al., Quantifying Dynamics in Phase-Separated Condensates Using Fluorescence Recovery after Photobleaching. Biophysical Journal, 2019. 117(7): p. 1285–1300.

75. Tinevez, J.-Y., et al., TrackMate: An open and extensible platform for single-particle tracking. Methods, 2017. 115: p. 80–90.

76. Ilavsky, J. and P.R. Jemian, Irena: tool suite for modeling and analysis of small-angle scattering. Journal of Applied Crystallography, 2009. 42(2): p. 347–353.

77. Thomas, K., et al., A multi-laboratory evaluation of a common in vitro pepsin digestion assay protocol used in assessing the safety of novel proteins. Regul Toxicol Pharmacol, 2004. 39(2): p. 87–98.

