## Supplementary material for "Intrinsically Disordered Protein Coating for Oral Delivery of Peptide Drugs": SI pdf

**Supplementary Information**

Table of Contents:


I. Table of Protein Sequences
II. Supplementary Figures
  a. Phase transition behaviors of proteins
  b. Material properties of condensates
  c. Glutamic Acid Mutations of SynIDP
  d. SynIDP Morphologies
  e. Peptide drug synthesis and encapsulation
  f. In vivo validation
  g. Mass Spectrometry of proteins/peptides

### I. Table S1 – Protein Amino Acid and DNA Sequences

| Protein Name | Amino Acid Sequence | DNA Sequence |
|---|---|---|
| synLCD | GVPGVGVPGVGVPGVGVPGV<br>GVPGVGVPGVGVPGVGVPGV<br>GVPGVGVPGVGVPGVGVPGV<br>GVPGVGVPGVGVPGVGTSTTE<br>TVAASAVAAVFEGPVPGVGVPG<br>VGVPGVGVPGVGVPGVGVPG<br>VGVPGVGVPGVGVPGVGVPG<br>VGVPGVGVPGVGVPGVGVPG<br>VGVPGVGTSTTETVAASAVAAV<br>FEGPVPGVGVPGVGVPGVGVP<br>GVGVPGVGVPGVGVPGVGVP<br>GVGVPGVGVPGVGVPGVGVP<br>GVGVPGVGVPGVGVPGVGVP<br>GVGVPGVGVPGVGVPGVGVP<br>GVGY | atgggggtaccgggcgtgggggttcctggagttggggtccct<br>ggagtaggtgtgccgggagtaggggtgcctggtgtcggggt<br>acctggcgtcggagtacctggggtaggagttccgggtgtgg<br>gtgttccaggcgttggtgtaccaggagtcggtgtcccaggag<br>tgggagtccccggtgtaggagtaccaggggtcggcgtacca<br>ggtgttggagtgccaggagtaggcacaagcacaacagaga<br>cagtcgcggcgtctgctgttgctgccgttttcgagggggtacc<br>gggcgtgggggttcctggagttggggtccctggagtaggtgt<br>gccgggagtaggggtgcctggtgtcggggtacctggcgtcg<br>gagtacctggggtaggagttccgggtgtgggtgttccaggcg<br>ttggtgtaccaggagtcggtgtcccaggagtgggagtccccg<br>gtgtaggagtaccaggggtcggcgtaccaggtgttggagtg<br>ccaggagtaggcacaagcacaacagagacagtcgcggcgt<br>ctgctgttgctgccgttttcgagggggtcccaggagttggtgtc<br>ccgggggttggagtacctggagtcggtgttcctggagtgggg<br>gtccctggagtgggtgttccgggcgtaggtgtcccaggtgtg<br>ggcgtaccgggcgttggtgttcctggtgtcggcgtgccgggt<br>gtgggtgttccgggcgtaggtgtcccaggtgtgggcgtaccg<br>ggcgttggtgttcctggtgtcggcgtgccgggtgtgggtgttc<br>cgggcgtaggtgtcccaggtgtgggcgtaccgggcgttggt<br>gttcctggtgtcggcgtgccgggctactgataatga |
| synLCD E1A | GVPGVGVPGVGVPGVGVPGV<br>GVPGVGVPGVGVPGVGVPGV<br>GVPGVGVPGVGVPGVGVPGV<br>GVPGVGVPGVGVPGVGTSTTA<br>TVAASAVAAVFEGPVPGVGVPG<br>VGVPGVGVPGVGVPGVGVPG<br>VGVPGVGVPGVGVPGVGVPG<br>VGVPGVGVPGVGVPGVGVPG<br>VGVPGVGTSTTATVAASAVAAV<br>FEGPVPGVGVPGVGVPGVGVP<br>GVGVPGVGVPGVGVPGVGVP<br>GVGVPGVGVPGVGVPGVGVP<br>GVGVPGVGVPGVGVPGVGVP<br>GVGVPGVGVPGVGVPGVGVP<br>GVGY | atgggggtaccgggcgtgggggttcntggagttggggtccc<br>tggagttggtgtgccgggagtaggggtgcctggtgtcggggt<br>acctggcgtcggagtacctggggtaggagttccgggtgtgg<br>gtgttccaggcgttggtgtaccaggagtcggtgtcccaggag<br>tgggagtccccggtgtaggagtaccaggggtcggcgtacca<br>ggtgttggagtgccaggagtaggcacctctactacggccacc<br>gtagcggcatcagctgtcgcggcagtattcgaaggggtacc<br>gggcgtgggggttcctggagttggggtccctggagtaggtgt<br>gccgggagtaggggtgcctggtgtcggggtacctggcgtcg<br>gagtacctggggtaggagttccgggtgtgggtgttccaggcg<br>ttggtgtaccaggagtcggtgtcccaggagtgggagtccccg<br>gtgtaggagtaccaggggtcggcgtaccaggtgttggagtg<br>ccaggagtaggcacctctactacggccaccgtagcggcatc<br>agctgtcgcggcagtattcgaaggggtcccaggagttggtgt<br>cccgggggttggagtacctggagtcggtgttcctggagtggg<br>ggtccctggagtgggtgttccgggcgtaggtgtcccaggtgt<br>gggcgtaccgggcgttggtgttcctggtgtcggcgtgccggg<br>tgtgggtgttccgggcgtaggtgtcccaggtgtgggcgtacc<br>gggcgttggtgttcctggtgtcggcgtgccgggtgtgggtgtt |

| | | |
|---|---|---|
| | | ccgggcgtaggtgtcccaggtgtgggcgtaccgggcgttgg<br>tgttcctggtgtcggcgtgccgggctactgataatga |
| synLCD<br>E2A | GVPGVGVPGVGVPGVGVPGV<br>GVPGVGVPGVGVPGVGVPGV<br>GVPGVGVPGVGVPGVGVPGV<br>GVPGVGVPGVGVPGVGTSTTE<br>TVAASAVAAVFAGPVPGVGVPG<br>VGVPGVGVPGVGVPGVGVPG<br>VGVPGVGVPGVGVPGVGVPG<br>VGVPGVGVPGVGVPGVGVPG<br>VGVPGVGTSTTETVAASAVAAV<br>FAGPVPGVGVPGVGVPGVGVP<br>GVGVPGVGVPGVGVPGVGVP<br>GVGVPGVGVPGVGVPGVGVP<br>GVGVPGVGVPGVGVPGVGVP<br>GVGVPGVGVPGVGVPGVGVP<br>GVGY | atgggggtaccgggcgtgggggttcctggagttggggtccct<br>ggagtaggtgtgccgggagtaggggtgcctggtgtcggggt<br>acctggcgtcggagtacctggggtaggagttccgggtgtgg<br>gtgttccaggcgttggtgtaccaggagtcggtgtcccaggag<br>tgggagtccccggtgtaggagtaccaggggtcggcgtacca<br>ggtgttggagtgccaggagtaggcacttcaaccactgagacc<br>gtagcagcctctgcggtggctgcagttttcgccggggtaccg<br>ggcgtgggggttcctggagttggggtccctggagtaggtgtg<br>ccgggagtaggggtgcctggtgtcggggtacctggcgtcgg<br>agtacctggggtaggagttccgggtgtgggtgttccaggcgtt<br>ggtgtaccaggagtcggtgtcccaggagtgggagtccccgg<br>tgtaggagtaccaggggtcggcgtaccaggtgttggagtgcc<br>aggagtaggcacttcaaccactgagaccgtagcagcctctgc<br>ggtggctgcagttttcgccggtgtcccaggagttggtgtcccg<br>ggggttggagtacctggagtcggtgttcctggagtgggggtc<br>cctggagtgggtgttccgggcgtaggtgtcccaggtgtgggc<br>gtaccgggcgttggtgttcctggtgtcggcgtgccgggtgtg<br>ggtgttccgggcgtaggtgtcccaggtgtgggcgtaccggg<br>cgttggtgttcctggtgtcggcgtgccgggtgtgggtgttccg<br>ggcgtaggtgtcccaggtgtgggcgtaccgggcgttggtgtt<br>cctggtgtcggcgtgccgggctactgataatga |
| ELP-<br>tev-<br>Exendin<br>4 | MGVGVPGVGVPGVGVPGVGV<br>PGVGVPGVGVPGVGVPGVGVP<br>GVGVPGVGVPGVGVPGVGVP<br>GVGVPGVGVPGVGVPGVGVP<br>GVGVPGVGVPGVGVPGVGVP<br>GVGVPGVGVPGVGVPGVGVP<br>GVGVPGVGVPGVGVPGVGVP<br>GVGVPGVGVPGVGVPGVGVP<br>GVGVPGVGVPGVGVPGHHHH<br>HHGVGVPGVGVPGVGVPGVG<br>VPGVGVPGVGVPGVGVPGVG<br>VPGVGVPGVGVPGVGVPGVG<br>VPGVGVPGVGVPGVGVPGVG<br>VPGVGVPGVGVPGVGVPGVG<br>VPGVGVPGVGVPGVGVPGVG<br>VPGVGVPGVGVPGVGVPGVG<br>VPGVGVPGVGVPGVGVPGVG<br>VPGVGVPGVGVPGVGVPGENL<br>YFQAAHGEGTFTSDLSKQMEE<br>EAVRLFIEWLKNGGPSSGAPPP<br>S | atgggtgtgggtgttccgggcgtaggtgtcccaggtgtgggc<br>gtaccgggcgttggtgttcctggtgtcggcgtgccgggtgtg<br>ggtgttccgggcgtaggtgtcccaggtgtgggcgtaccggg<br>cgttggtgttcctggtgtcggcgtgccgggtgtgggtgttccg<br>ggcgtaggtgtcccaggtgtgggcgtaccgggcgttggtgtt<br>cctggtgtcggcgtgccgggtgtgggtgttccgggcgtaggt<br>gtcccaggtgtgggcgtaccgggcgttggtgttcctggtgtcg<br>gcgtgccgggtgtgggtgttccgggcgtaggtgtcccaggtg<br>tgggcgtaccgggcgttggtgttcctggtgtcggcgtgccgg<br>gtgtgggtgttccgggcgtaggtgtcccaggtgtgggcgtac<br>cgggcgttggtgttcctggtgtcggcgtgccgggtgtgggtgt<br>tccgggcgtaggtgtcccaggtgtgggcgtaccgggcgttg<br>gtgttcctggtgtcggcgtgccgggccatcaccatcaccatca<br>tggtgtgggtgttccgggcgtaggtgtcccaggtgtgggcgt<br>accgggcgttggtgttcctggtgtcggcgtgccgggtgtggg<br>tgttccgggcgtaggtgtcccaggtgtgggcgtaccgggcgt<br>tggtgttcctggtgtcggcgtgccgggtgtgggtgttccgggc<br>gtaggtgtcccaggtgtgggcgtaccgggcgttggtgttcctg<br>gtgtcggcgtgccgggtgtgggtgttccgggcgtaggtgtcc<br>caggtgtgggcgtaccgggcgttggtgttcctggtgtcggcg<br>tgccgggtgtgggtgttccgggcgtaggtgtcccaggtgtgg |

| | | |
|---|---|---|
| | | gcgtaccgggcgttggtgttcctggtgtcggcgtgccgggtg<br>tgggtgttccgggcgtaggtgtcccaggtgtgggcgtaccgg<br>gcgttggtgttcctggtgtcggcgtgccgggtgtgggtgttcc<br>gggcgtaggtgtcccaggtgtgggcgtaccgggcgttggtgt<br>tcctggtgtcggcgtgccgggcgagaatttatatttccaagcc<br>gcgcatggggagggcacatttacatccgatcttagcaagcag<br>atggaggaagaggcggtgcgtctgtttattgaatggttaaaga<br>acggaggacccagttccggcgctccgcctccctct |
| Exendin 4 | AAHGEGTFTSDLSKQMEEEAV<br>RLFIEWLKNGGPSSGAPPPSG | |
| ELP-tev-ExCDC | MGVGVPGVGVPGVGVPGVGV<br>PGVGVPGVGVPGVGVPGVGVP<br>GVGVPGVGVPGVGVPGVGVP<br>GVGVPGVGVPGVGVPGVGVP<br>GVGVPGVGVPGVGVPGVGVP<br>GVGVPGVGVPGVGVPGVGVP<br>GVGVPGVGVPGVGVPGVGVP<br>GVGVPGVGVPGVGVPGVGVP<br>GVGVPGVGVPGVGVPGHHHH<br>HHGVGVPGVGVPGVGVPGVG<br>VPGVGVPGVGVPGVGVPGVG<br>VPGVGVPGVGVPGVGVPGVG<br>VPGVGVPGVGVPGVGVPGVG<br>VPGVGVPGVGVPGVGVPGVG<br>VPGVGVPGVGVPGVGVPGVG<br>VPGVGVPGVGVPGVGVPGVG<br>VPGVGVPGVGVPGVGVPGVG<br>VPGVGVPGVGVPGVGVPGENL<br>YFQAAHGEGTFTSDLSKQMEE<br>EAVRLFIEWLKNGGPSSGAPPP<br>SGTSTTETVAASAVAAVFEG | atgggtgtgggtgttccgggcgtaggtgtcccaggtgtgggc<br>gtaccgggcgttggtgttcctggtgtcggcgtgccgggtgtg<br>ggtgttccgggcgtaggtgtcccaggtgtgggcgtaccggg<br>cgttggtgttcctggtgtcggcgtgccgggtgtgggtgttccg<br>ggcgtaggtgtcccaggtgtgggcgtaccgggcgttggtgtt<br>cctggtgtcggcgtgccgggtgtgggtgttccgggcgtaggt<br>gtcccaggtgtgggcgtaccgggcgttggtgttcctggtgtcg<br>gcgtgccgggtgtgggtgttccgggcgtaggtgtcccaggtg<br>tgggcgtaccgggcgttggtgttcctggtgtcggcgtgccgg<br>gtgtgggtgttccgggcgtaggtgtcccaggtgtgggcgtac<br>cgggcgttggtgttcctggtgtcggcgtgccgggtgtgggtgt<br>tccgggcgtaggtgtcccaggtgtgggcgtaccgggcgttg<br>gtgttcctggtgtcggcgtgccgggccatcaccatcaccatca<br>tggtgtgggtgttccgggcgtaggtgtcccaggtgtgggcgt<br>accgggcgttggtgttcctggtgtcggcgtgccgggtgtggg<br>tgttccgggcgtaggtgtcccaggtgtgggcgtaccgggcgt<br>tggtgttcctggtgtcggcgtgccgggtgtgggtgttccgggc<br>gtaggtgtcccaggtgtgggcgtaccgggcgttggtgttcctg<br>gtgtcggcgtgccgggtgtgggtgttccgggcgtaggtgtcc<br>caggtgtgggcgtaccgggcgttggtgttcctggtgtcggcg<br>tgccgggtgtgggtgttccgggcgtaggtgtcccaggtgtgg<br>gcgtaccgggcgttggtgttcctggtgtcggcgtgccgggtg<br>tgggtgttccgggcgtaggtgtcccaggtgtgggcgtaccgg<br>gcgttggtgttcctggtgtcggcgtgccgggtgtgggtgttcc<br>gggcgtaggtgtcccaggtgtgggcgtaccgggcgttggtgt<br>tcctggtgtcggcgtgccgggcgagaatttatatttccaagcc<br>gcgcatggggagggcacatttacatccgatcttagcaagcag<br>atggaggaagaggcggtgcgtctgtttattgaatggttaaaga<br>acggaggacccagttccggcgctccgcctccctctgggacat<br>cgacaactgagaccgtcgctgccagtgctgtggcagcagtgt<br>ttgagggctgataatga |
| ExCDC | AAHGEGTFTSDLSKQMEEEAV<br>RLFIEWLKNGGPSSGAPPPSGTS<br>TTETVAASAVAAVFEG | |

### II. Supplementary Figures

#### a. Phase transitions

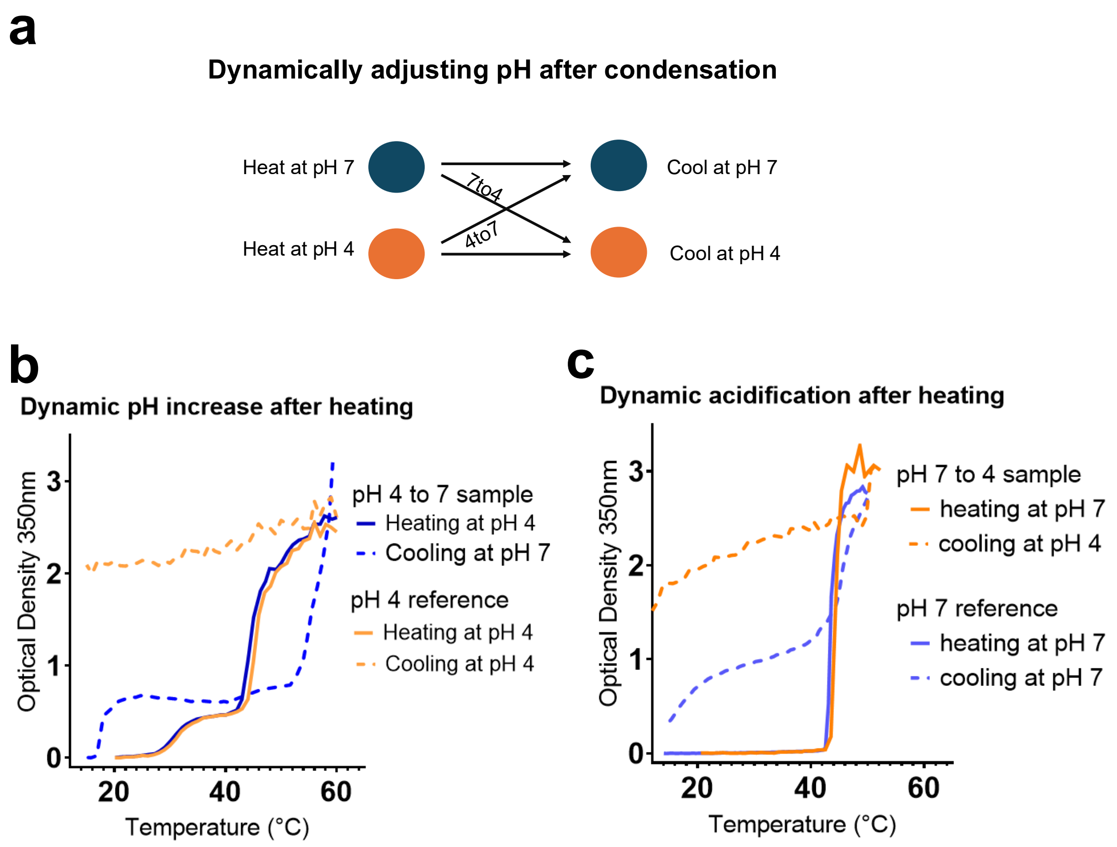


**Figure S1:** Adjusting the pH of heated condensate solutions. **a,** A schematic of the four different conditions tested in which heating occurs at pH 4 or 7 in 10mM citrate-phosphate buffer, then pH was adjusted from 4 to 7 or from 7 to 4 in dynamic samples. Unchanged samples were also included for reference. **b,** Adjusting heated pH 4 samples to pH 7 resulted in reversible cooling profiles, as opposed to an irreversible profile at pH 4. **c,** Samples that were heated at pH 7 showed diverging reversibility profiles when cooled at pH 4 or pH 7.

#### b. Material properties of condensates

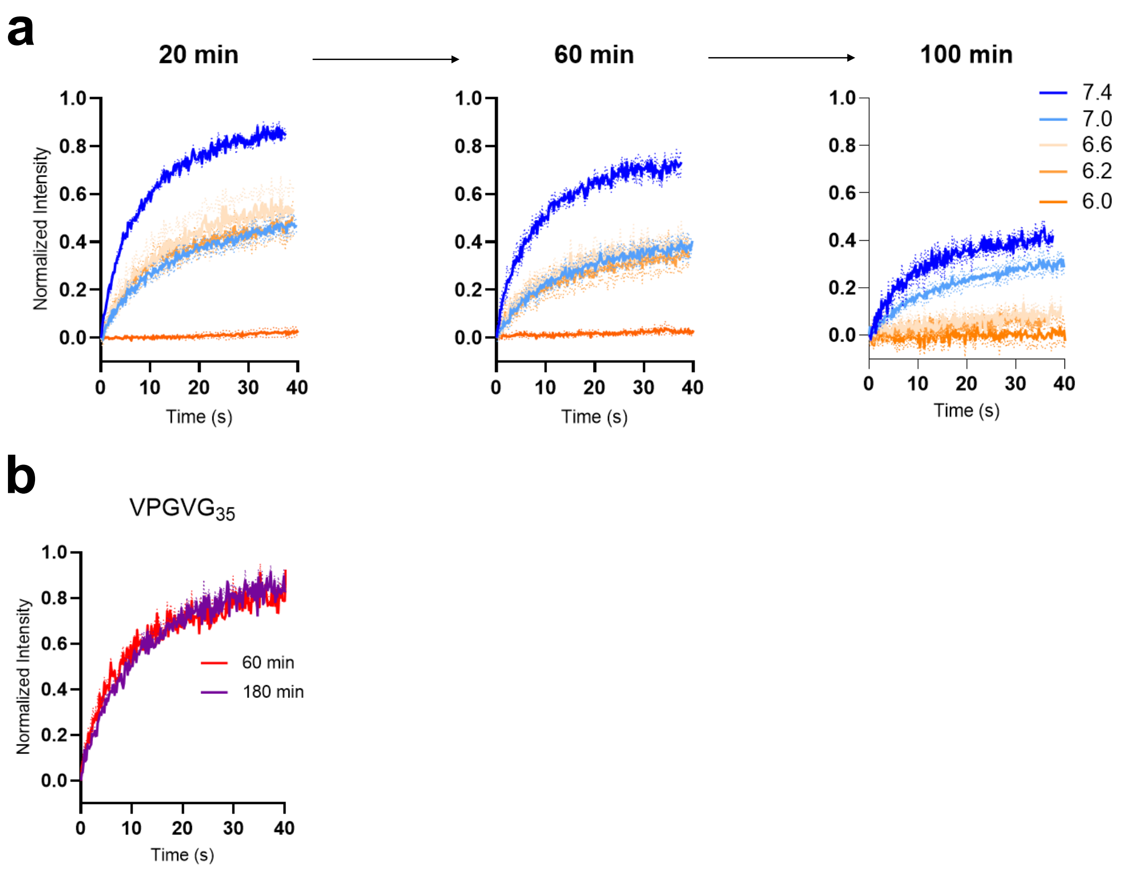


**Figure S2:** Additional FRAP recovery curves of aging SynIDP samples at intermediate pHs and an ELP control. **a,** Recovery curves of samples between pH 6 and 7.4 tended to cluster at intermediate values between 30% and 60% recovery, while aging from 20 to 60 or 100 minutes resulted in reduced recovery over time. Samples at pH 6.2 and 6.2 reached near-zero recovery by 100 min. **b,** ELP condensate samples composed of 35 repeats of VPGVG recovered similar intensity to pH 7.4 SynIDP at early time points, and did not exhibit aging behavior up to 180 minutes.

**Span (fractional recovery)**

| pH | time (min) 20 | 60 | 100 | 180 |
|---|---|---|---|---|
| 6 | *-0.0286 | | | |
| 6.2 | 0.5342 | 0.2831 | *0.1309 | |
| 6.4 | 0.4515 | 0.3176 | *0.1343 | |
| 6.6 | 0.5388 | 0.3643 | *0.1969 | *0.02349 |
| 6.8 | 0.4703 | 0.3946 | 0.2809 | |
| 7 | 0.4684 | 0.3777 | 0.3137 | |
| 7.2 | | 0.4656 | 0.3798 | |
| 7.4 | 0.7234 | 0.6623 | 0.4017 | 0.3194 |
| 8 | 0.7708 | 0.7425 | | 0.4369 |
| ELP | | 0.7238 | | 0.8264 |

**Table S2** – Maximum recovery values of additional pHs and times for SynIDP and ELP, derived from the exponential fitting of experimental curves in GraphPad Prism. Values with an asterisk denote poor fitting to a 1-phase association model, with R squared values below 0.75. Colors correspond to the amount of fraction recovery, with red indicating low recovery and blue indicating high recovery.

**Tau (recovery half-life)**

| pH \ time | 20 | 60 | 100 | 180 |
|---|---|---|---|---|
| 6 | *33.74 | n/a | n/a | n/a |
| 6.2 | 10.88 | 11.16 | *44.87 | |
| 6.4 | 10.47 | 12.69 | *12.13 | |
| 6.6 | 12.42 | 10.89 | *80.21 | n/a |
| 6.8 | 12.02 | 15.17 | 15.71 | |
| 7 | 13.77 | 12.93 | 15.92 | |
| 7.2 | | 10.1 | 11.33 | |
| 7.4 | 11.72 | 8.619 | 9.427 | 11.82 |
| 8 | 10.51 | 10.79 | | 13.18 |
| ELP | | 9.816 | | 12.23 |

**Table S3** – Speed of recovery for SynIDP and ELP recovery curves were also calculated from exponential fitting. Exponential fitting as a 1-phase process assumes a form of

$y = (span)(1 - \exp\,(-{}^{t}/_{tau})$, in which tau is a related to the halving time, in seconds. Most fully arrested recovery curves were unable to be fitted and are denoted as n/a in the table, while tau values from poorly fitted curves are marked with an asterisk. All curves that showed good fitting had similar tau values, ranging from ~8 sec to ~16 sec. Values are color coded with blue indicating fast recovery and orange indicating slow recovery times.

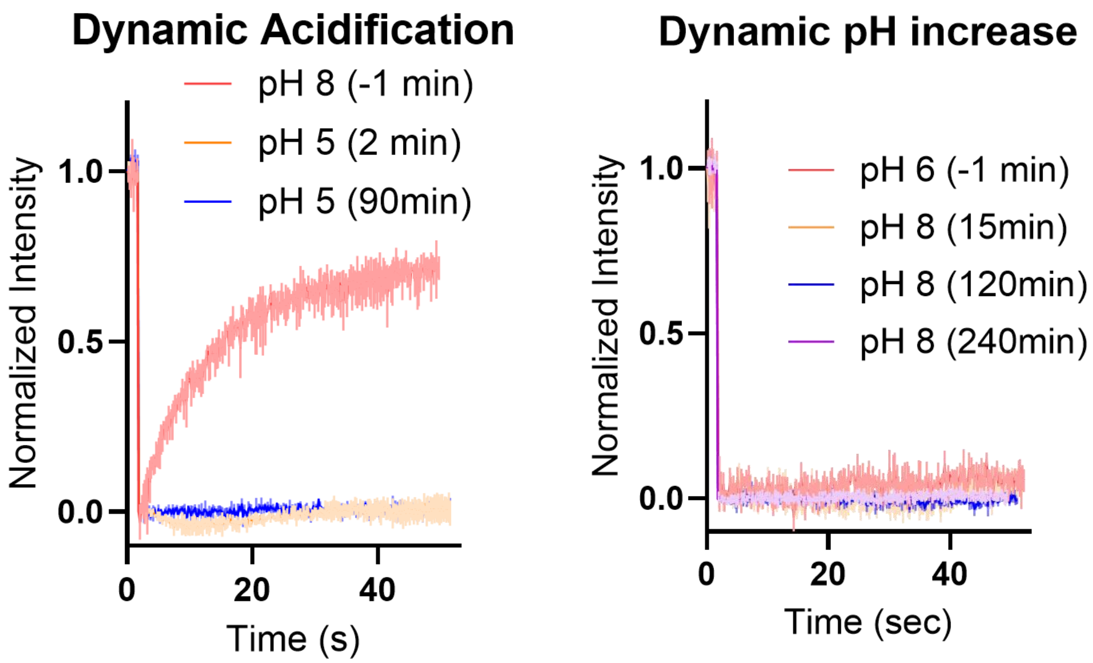


**Figure S3**: FRAP of condensates before and after pH changes. **left,** Dynamic acidification of liquid SynIDP condensates formed at pH 8 and aged for 1 hour results in immediate and persistent dynamical arrest. **right,** The opposite, adding base to arrested condensates to change the solution pH from 6 to 8, does not re-fluidize the samples, even after 6 hours.

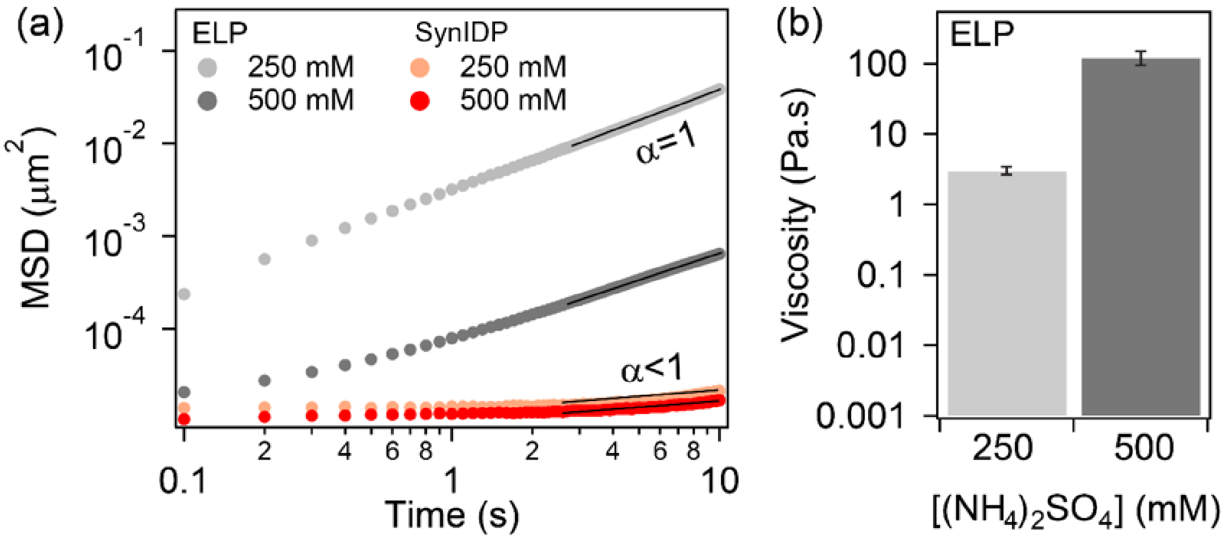


**Figure S4:** Salt dependent material properties of ELP and synLCD condensates, as measured by nanorheology. **a,** Ensemble-averaged mean-squared displacement of 200 nm beads embedded inside the ELP and synLCD condensates at 250 mM (ELP, light gray; SynIDP, orange) and 500 mM (ELP, dark gray; SynIDP, red) $(NH_4)_2SO_4$. **b,** Viscosity of ELP condensates determined from the MSDs shown in (a) at 250 mM (light gray) and 500 mM (dark gray) $(NH_4)_2SO_4$. The viscosity for synLCD condensates could not be determined due to the subdiffusive motion of the beads inside the synLCD condensates. All measurements have been replicated independently at least n = 3 times.

#### c. Glutamic acid mutations of SynIDP

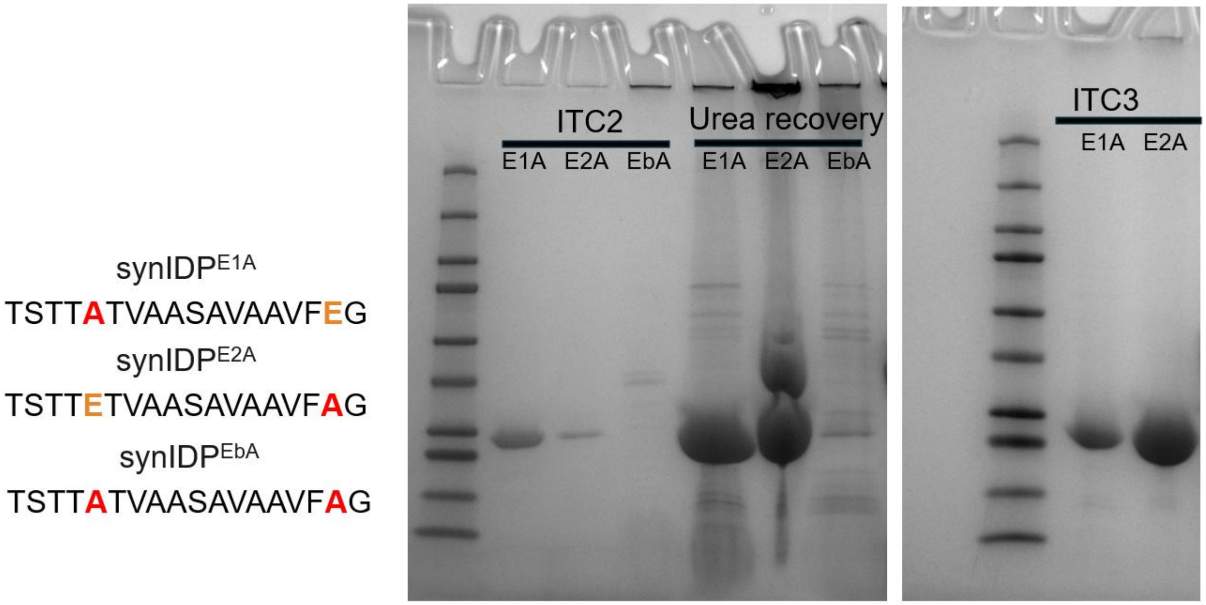


**Figure S5:** Mutating the glutamic acids of the CDC19 RAC in SynIDPs decreases protein yields from bacterial expression and purification. SDS-PAGE gels of the second round of ITC shows minimal protein from E1A or E2A soluble fractions, and none at the correct size for EbA. Both single mutants required additional solubilization in 2M urea for the third round of ITC, although the double mutant was intractable under any conditions. The “wild type” SynIDP did not require any additional solubilization steps, being readily reversible in cold, slightly alkaline buffers.

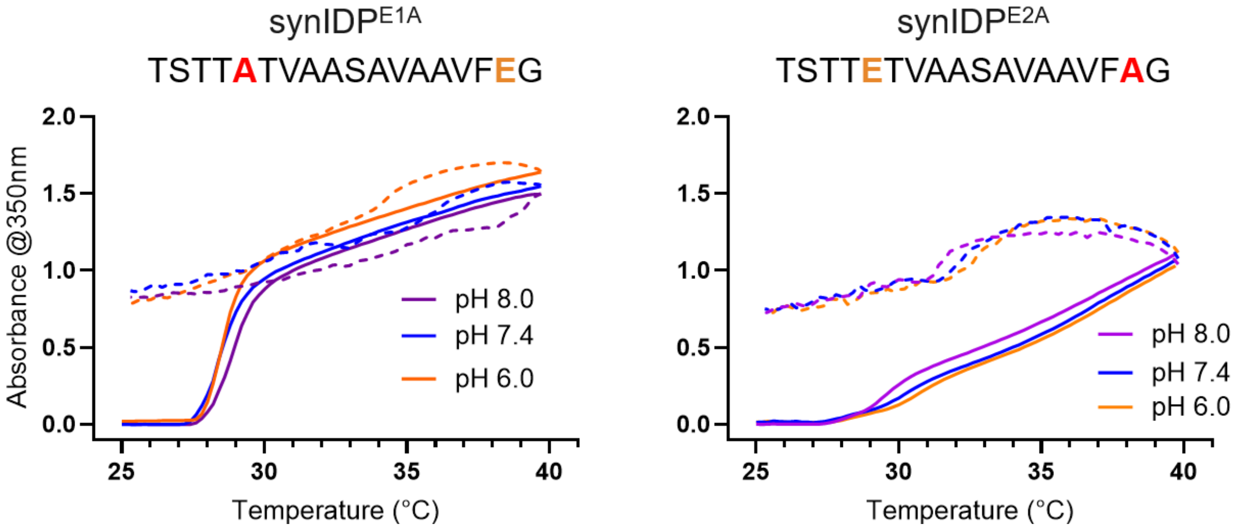


**Figure S6:** Consistently irreversible Cary UV-Vis curves of EtoA mutants. **left,** The E1A mutant in PBS at pH 6, 7.4, or 8 showed a lower transition temperature than SynIDP, and irreversible cooling curves at all pHs. **right,** Mutant E2A was similarly irreversible at all tested pHs, and uniquely exhibited a linear transition with heating instead of a sigmoidal transition curve.

#### d. SynIDP Morphologies

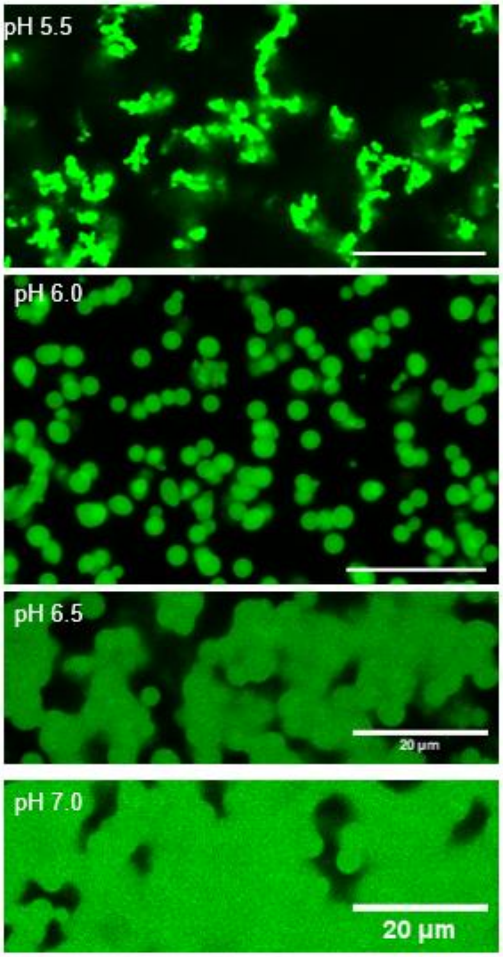


**Figure S7:** Morphologies generated by heating SynIDP at additional pHs. If the liquid-to-solid transition occurs faster than gravity-induced settling, as it does at pH 5.5, arrested globules stick together. At pH 6 the liquid-to-solid transition is slower, allowing larger droplet features to form before arrest. Higher pHs such as 6.5 and 7 feature puddle-like morphologies due to their more persistent viscoelastic behavior that exceeds the rate of droplet settling. All scale bars are 20µm.

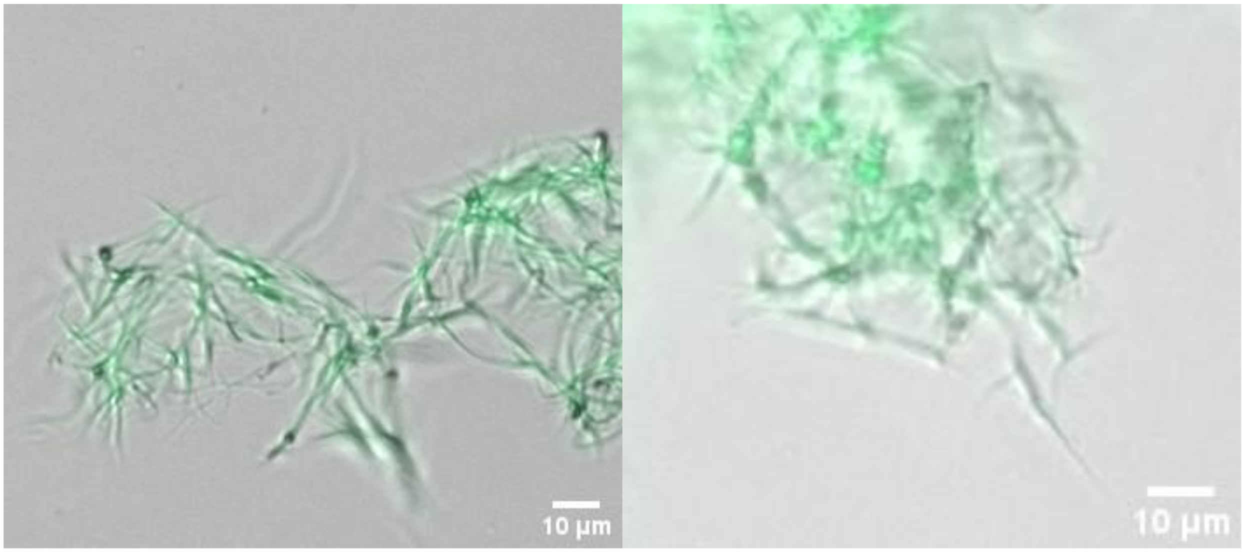


**Figure S8:** Superimposed bright field and ThT fluorescence imaging of SynIDP after incubation at pH 6 for 3 days at room temperature, then heating to 45 °C. Filamentous assemblies that are ThT positive and tens of microns in length form tangled networks.

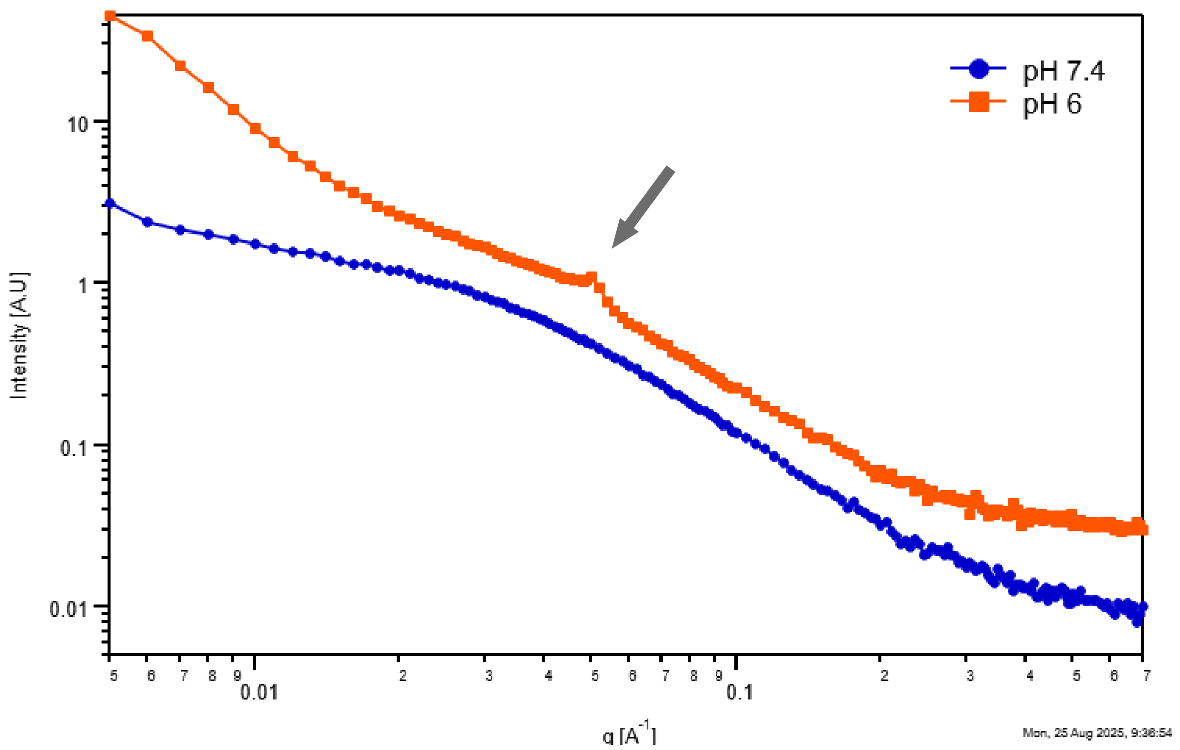


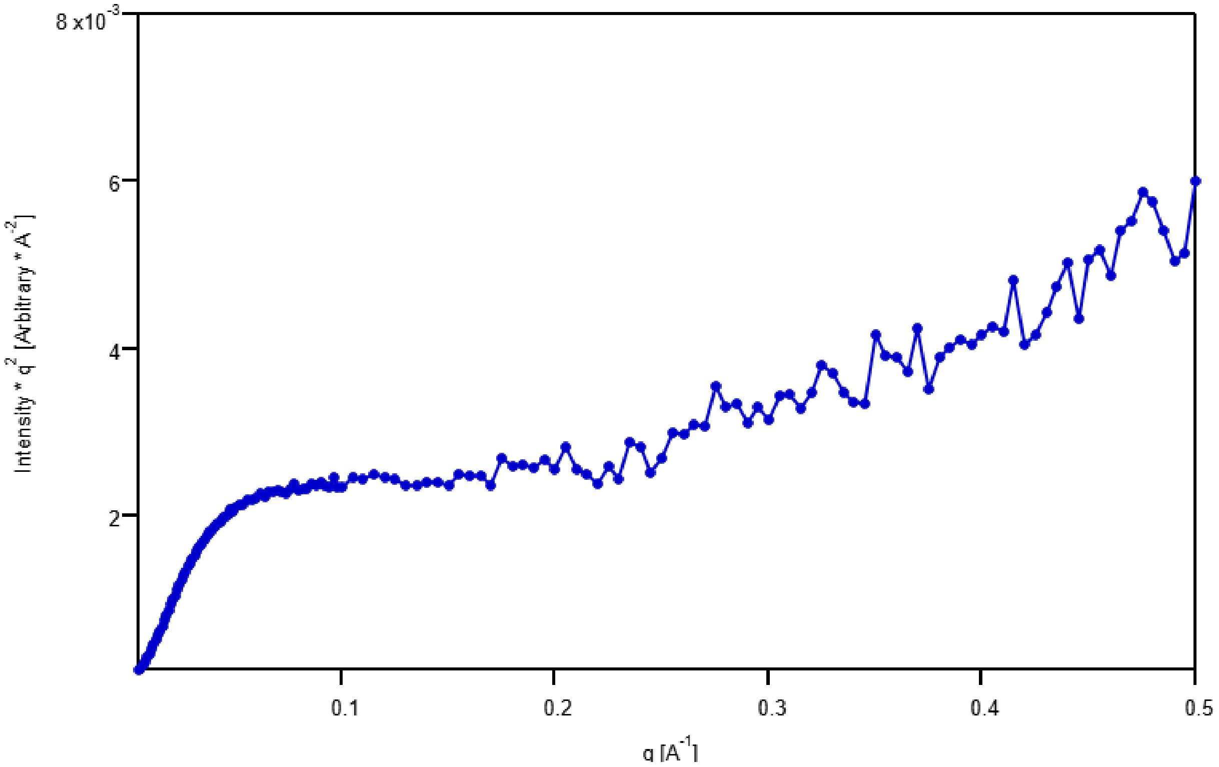


**Figure S9:** Small angle X-ray Scattering of synLCD at pH 6.0 or pH 7.4 at 25 °C shows a Bragg peak at pH 6.0 and no peak at pH 7.4 (top). Kratky Plot of SynLCD at pH 7.4 (bottom).

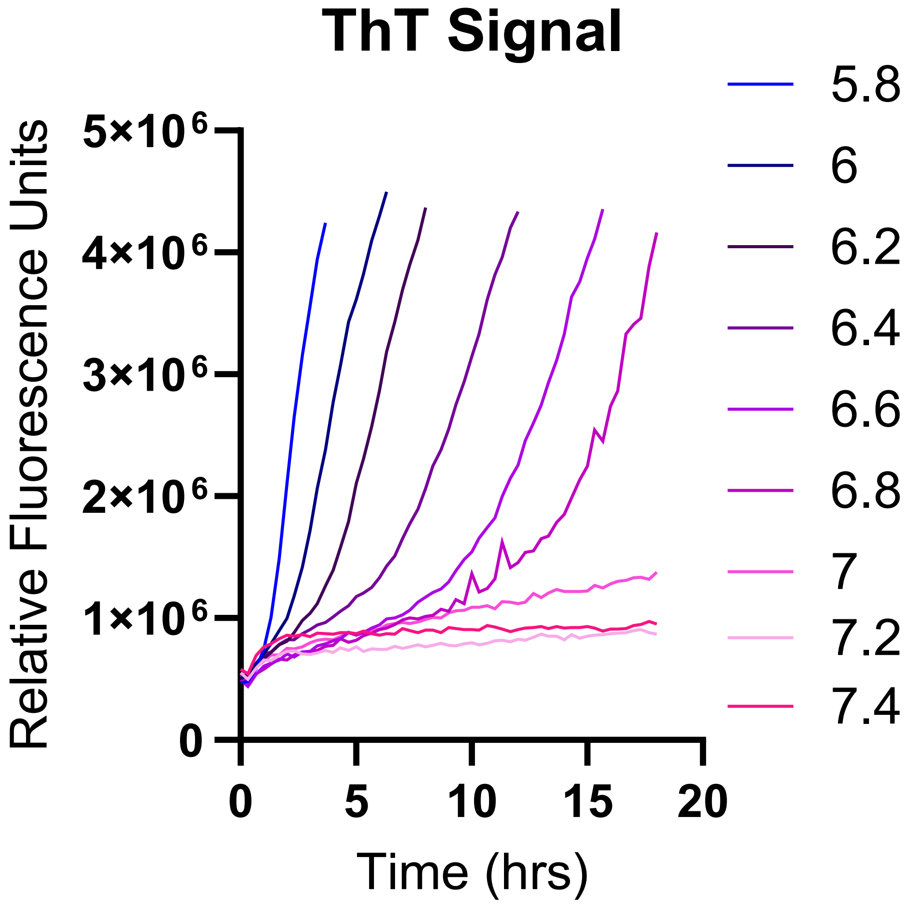


**Figure S10:** Monitoring fiber/filament growth at various pHs using ThT fluorescence over time in an automated plate reader (Spectramax iD5) with incubation at 37 °C.

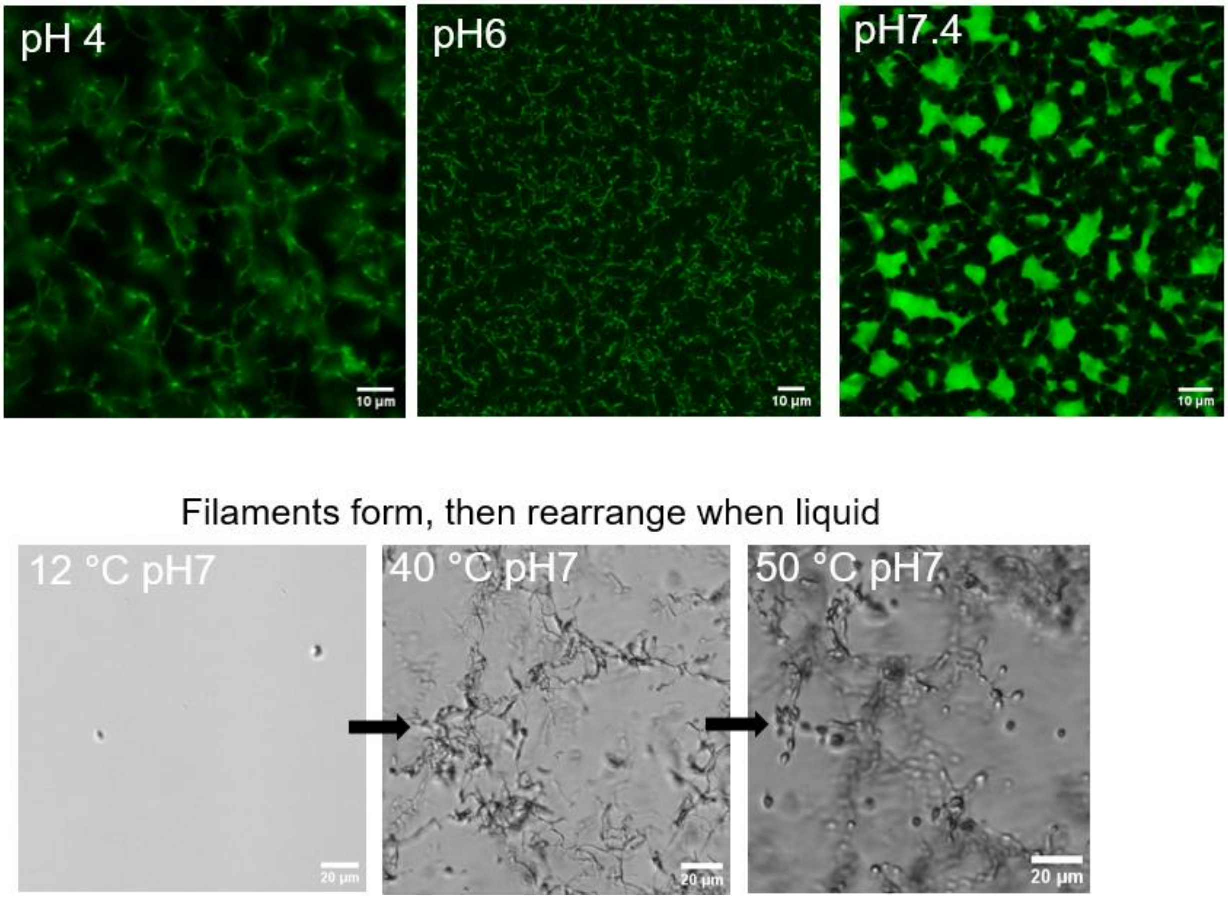


**Fig S11:** Manipulation of filament structures by modulating pH of nanofiber suspensions before thermal transitions (top), or modulating temperature after the initial thermal transition (bottom).

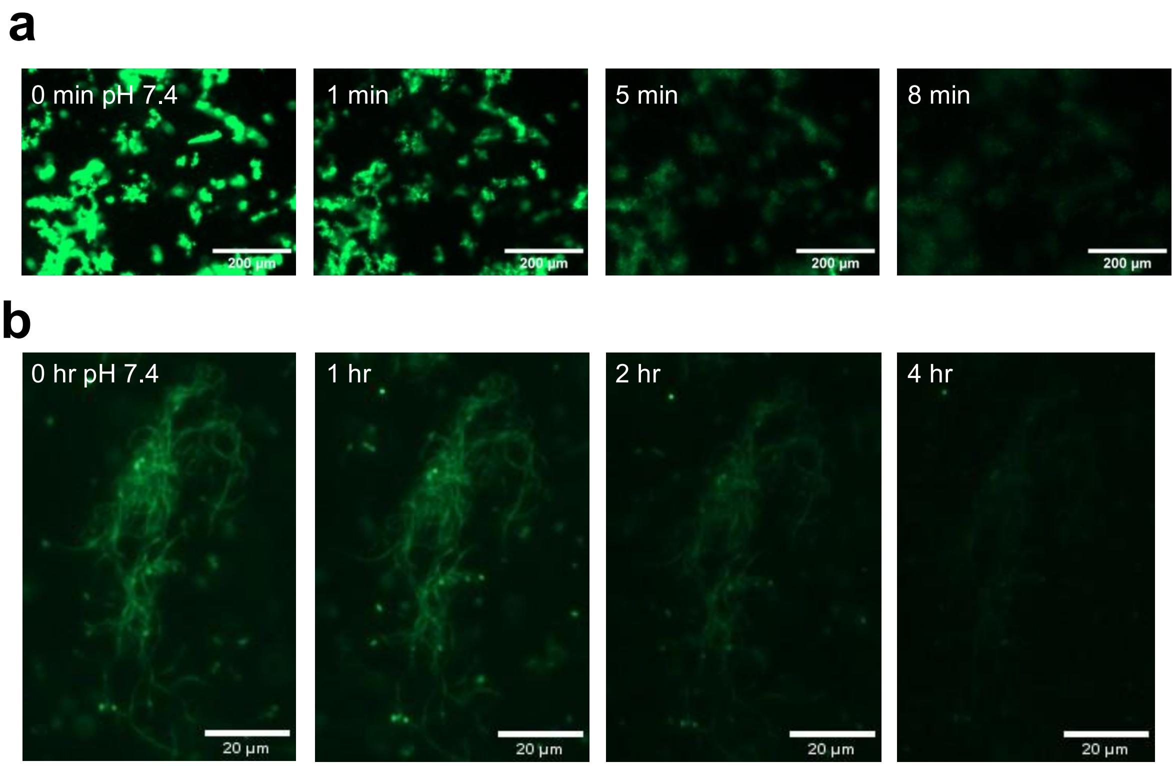


**Figure S12:** The dissolution of filament bundles and fibrils. **a,** Filamentous bundles formed at pH 6 were adjusted to pH 7.4 in PBS at 30 °C, then monitored by ThT fluorescence with 2 sec exposure times as they disperse. **b,** An individual fibril tangle is monitored periodically as it gradually dissolves.

#### e. Peptide drug synthesis and encapsulation

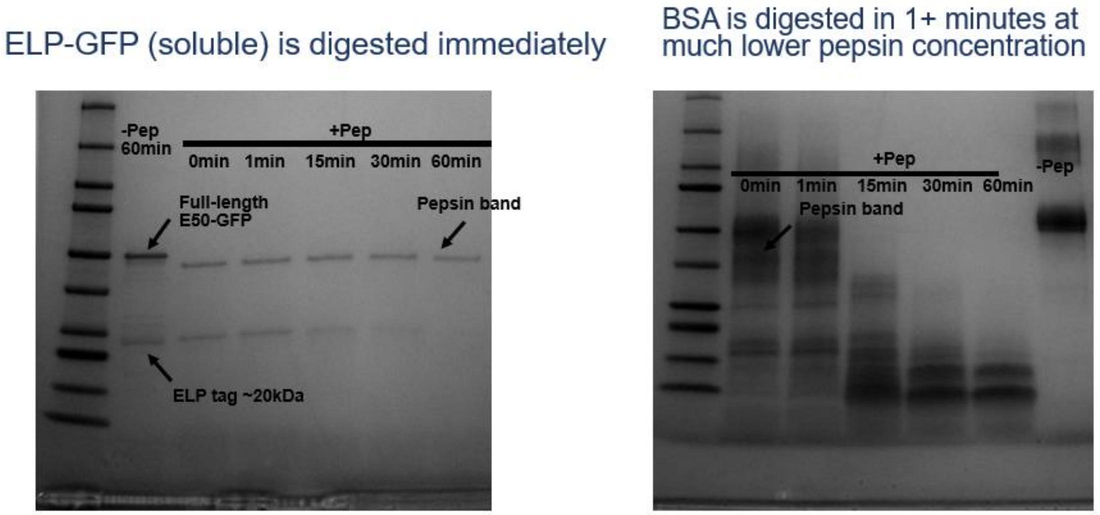


**Figure S13:** SDS-PAGE gels of proteins with more typical distributions of amino acids in SGA. **left,** A soluble ELP-GFP fusion is immediately digested, leaving the ELP tag that disappears after 15 minutes. **right,** Bovine serum albumin is digested rapidly even at very high BSA concentration and low pepsin concentration.

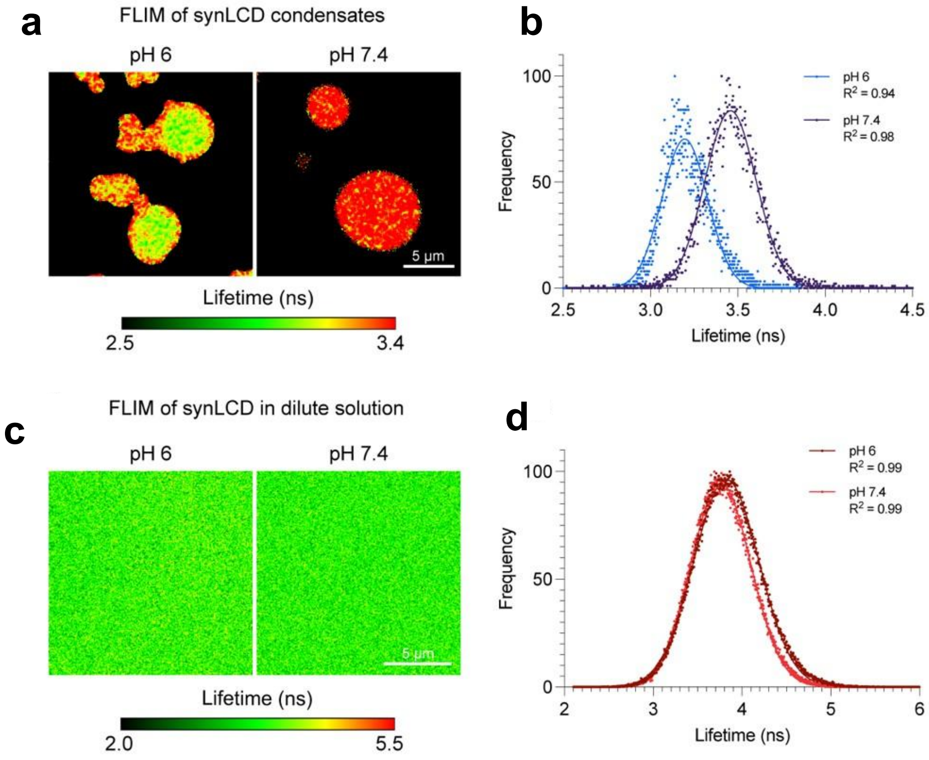


**Figure S14: a,** Frequency-domain fluorescence lifetime microscopy (FD-FLIM) measurements of synLCD condensates at two distinct pH conditions. **b,** Corresponding fluorescence lifetime distributions. **c,** FD-FLIM measurements of synLCD in dilute solution at two distinct pH conditions and **d,** corresponding fluorescence lifetime distributions.

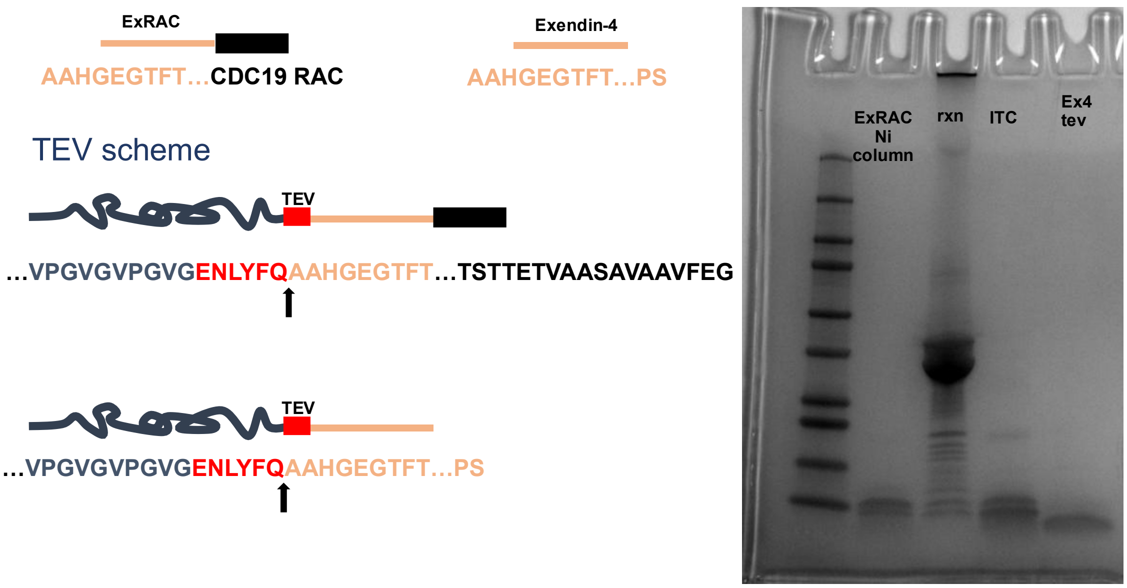


**Figure S15:** Synthesis scheme (left) for TEV-cleavable fusions of peptide drugs to ELP for purification, followed by cleavage, reverse ITC, and reverse NTA column steps to isolate pure peptide drugs Exendin-4 or ExRAC. Intermediate steps and pure peptides are analyzed by SDS-PAGE (right) for purity, and molecular weight is confirmed by MALDI-MS (**See Fig. S23**).

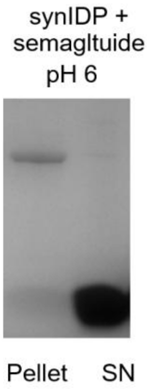


**Figure S16:** Encapsulation of semaglutide with SynIDP at pH 6. A semaglutide sample was dissolved in neutral citrate-phosphate buffer, then adjusted to pH 6. SynIDP was added, then heated to trigger encapsulation. The supernatant after centrifugation (right) is compared to the resuspended pellet (left) to show encapsulation efficiency.

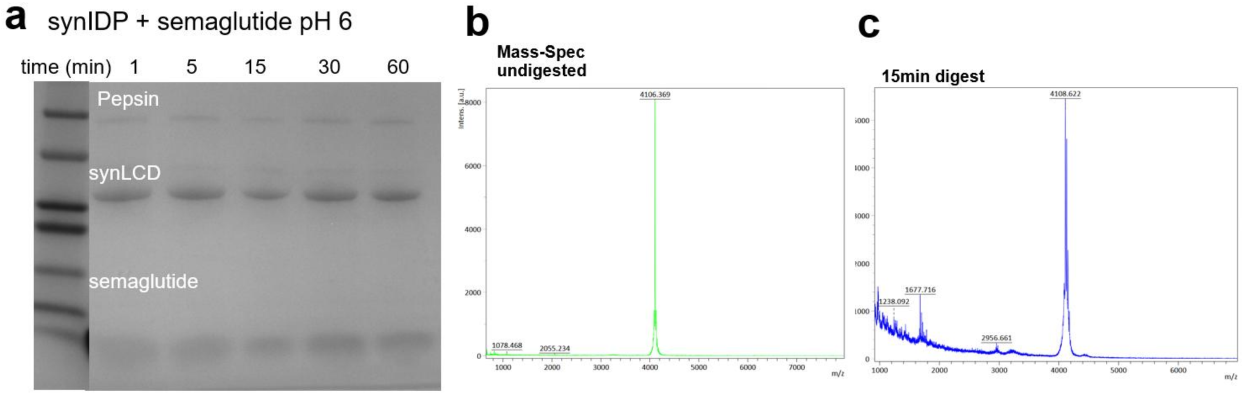


**Figure S17** – SynIDP protection of semaglutide. **a,** SDS-PAGE of encapsulated semaglutide in SynIDP. Semaglutide does not run at its predicted molecular weight due to the alkyl pendant, so we confirmed survival with MALDI-MS by comparing the undigested peak (**b**) to the peak after 15 min of SGA conditions (**c**).

**f. In-vivo validation**

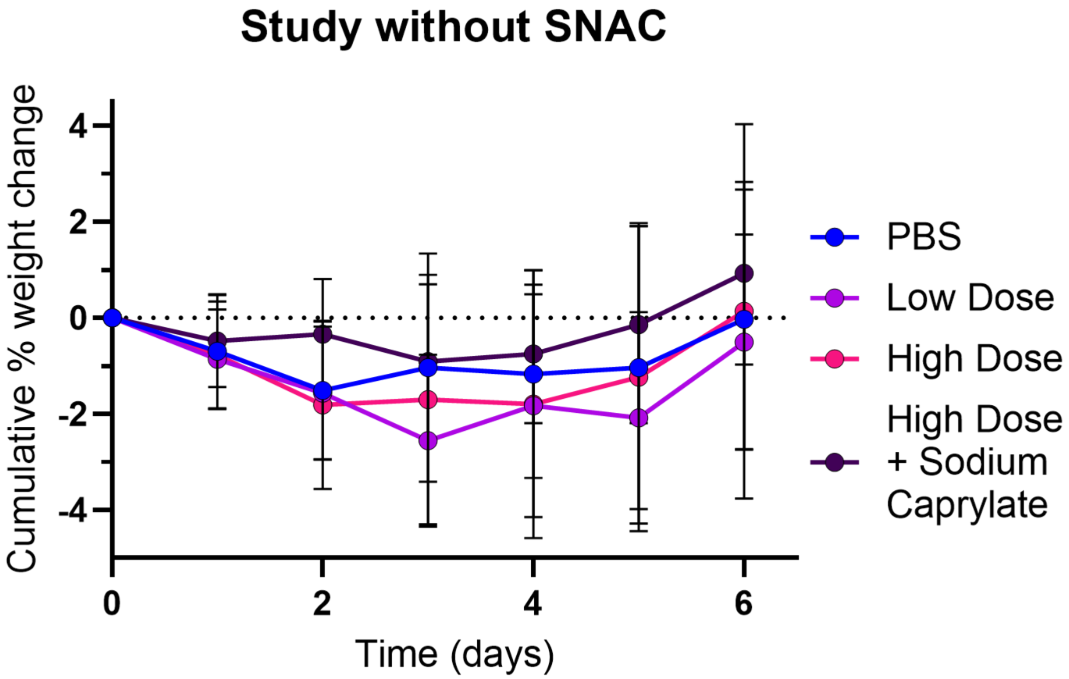


**Figure S18:** Oral delivery of peptide drugs without SNAC is ineffective. Normalized weights to the study start date for PBS, low ExCDC dosing (1000nm/kg), high ExCDC dosing (4000nm/kg), and high dosing with sodium caprylate (30mg/kg) show no significant weight changes. All groups are n=4.

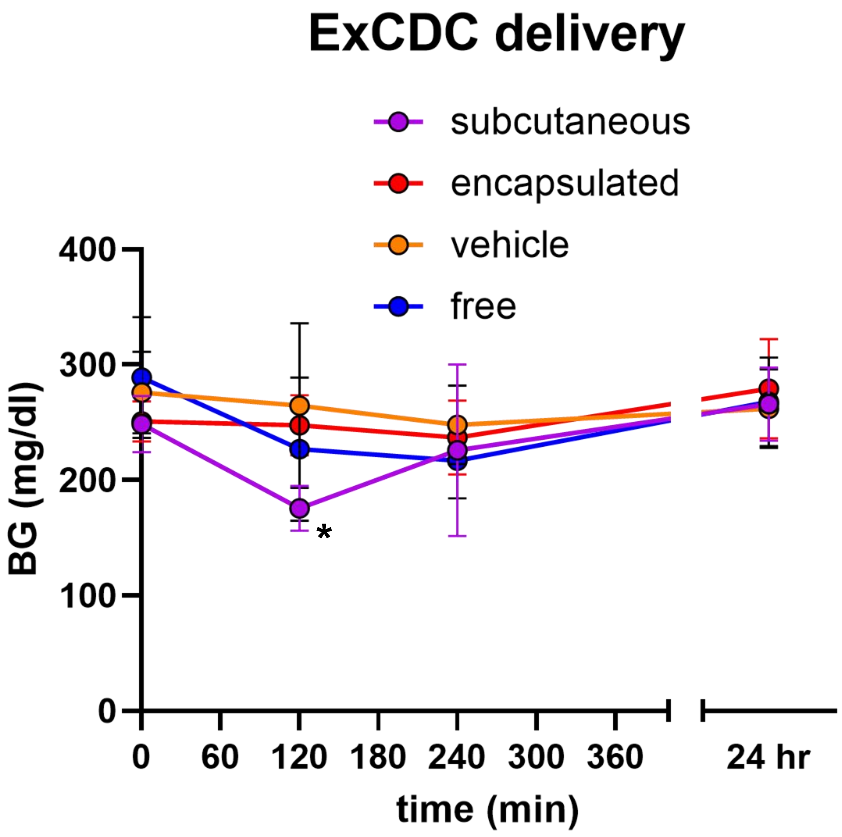


**Figure S19:** Blood glucose measurements after treatment with ExCDC or controls. Delivery of ExCDC by subcutaneous injection of free peptide (n=5) was significant against the oral vehicle group (n=5) at 120 min. Oral administration of encapsulated (n=7) or free ExCDC (n=7) with SNAC did not show significance against any groups.

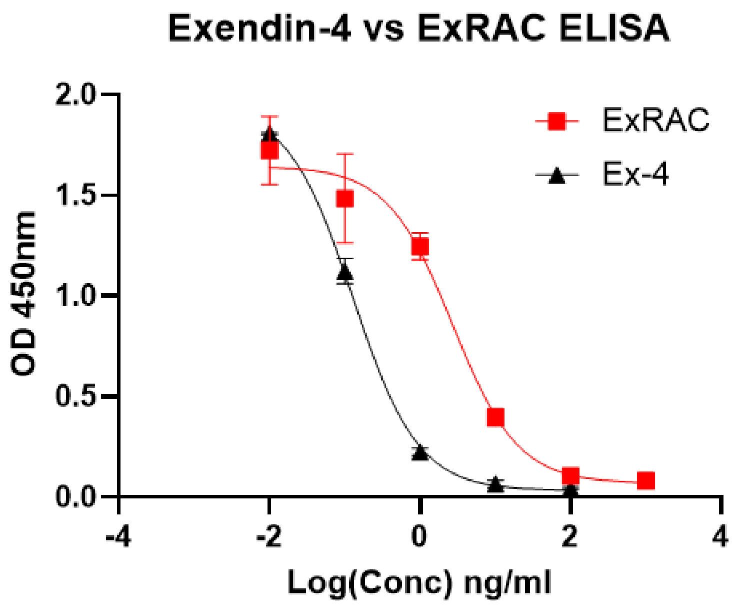


**Figure S20:** Competitive ELISA curves of Exendin-4 antibody binding to ExCDC doped into mouse serum versus the Exendin-4 control.

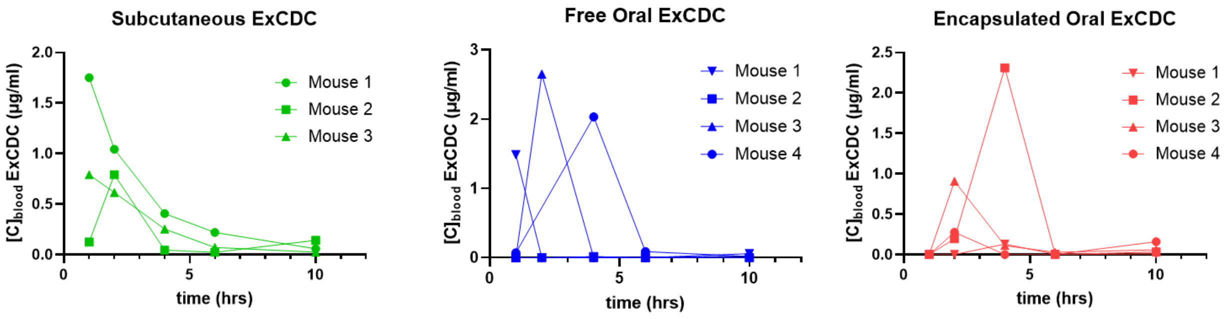


**Figure S21:** Serum was collected after administration and analyzed by ELISA for concentration of molecules binding to Exendin-4 antibodies, then blood levels were calculated using a standard curve generated for ExCDC in serum (see Supplementary Fig. 19). Blood levels from each mouse are plotted separately due to high variability.

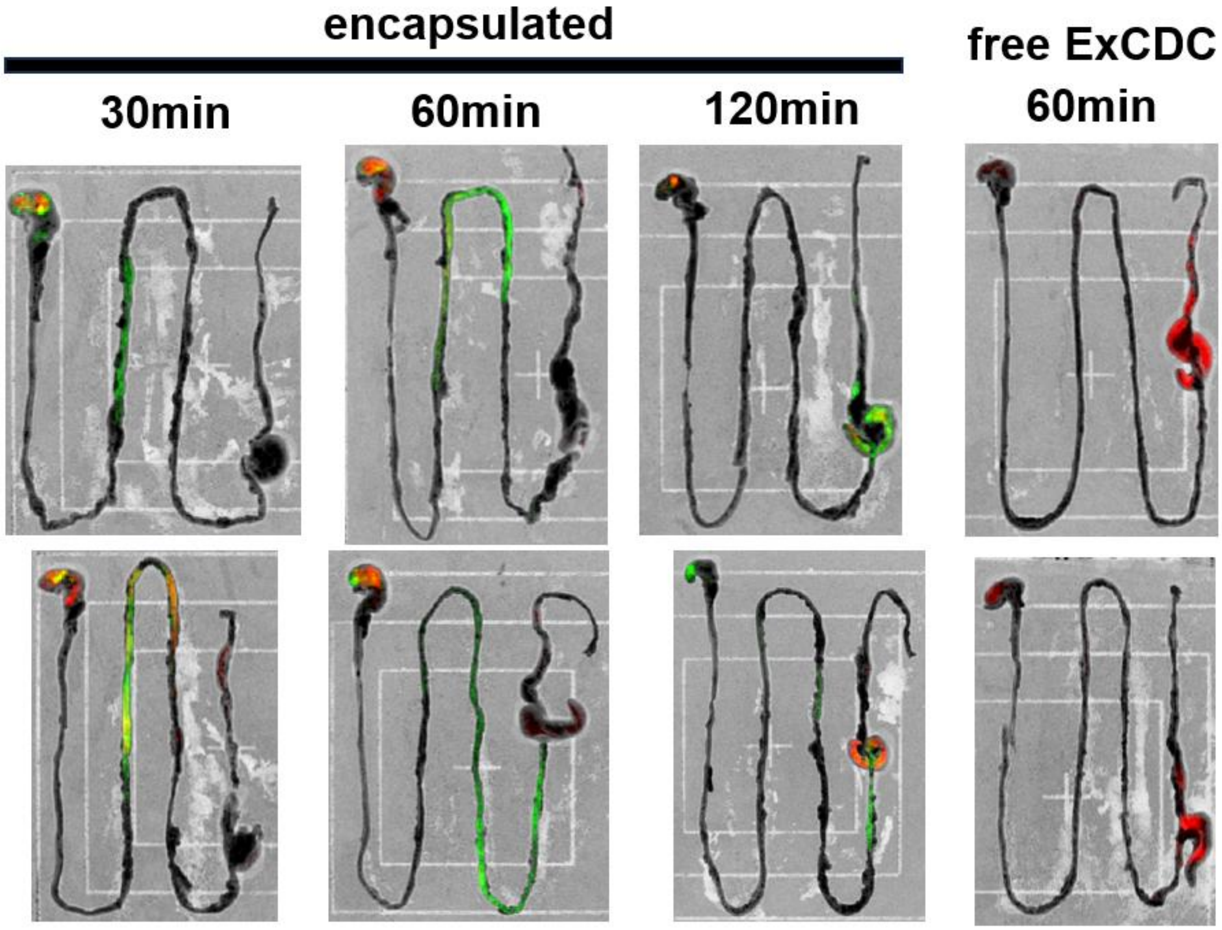


**Figure S22:** Superimposed fluorescence and brightfield images of dissected mouse GIs following oral administration of synIDP-encapsulated or free ExCDC at various timepoints. ExCDC is denoted in the red channel, SynIDP is denoted in the green channel. The outline of the GI from stomach on the upper left to cecum and lower GI on the right can be seen in the inverted brightfield images.

#### g. Mass-Spec of proteins/peptides

**Figure S23:** Mass spectroscopy confirmation of peptide and protein molecular weights.

Exendin-4: Predicted MW 4386 Da

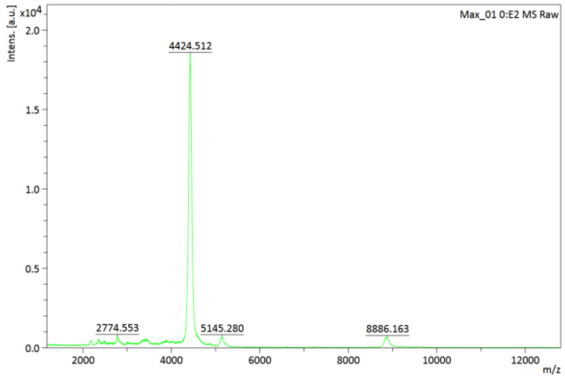


ExRAC: Predicted MW 6080 Da

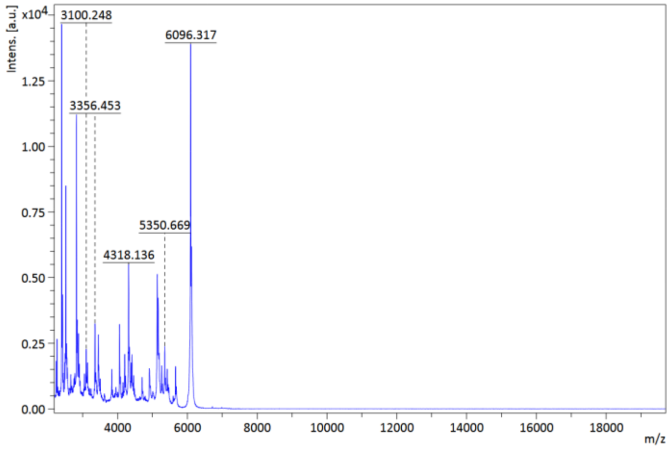


SynIDP: Predicted MW 24178 Da

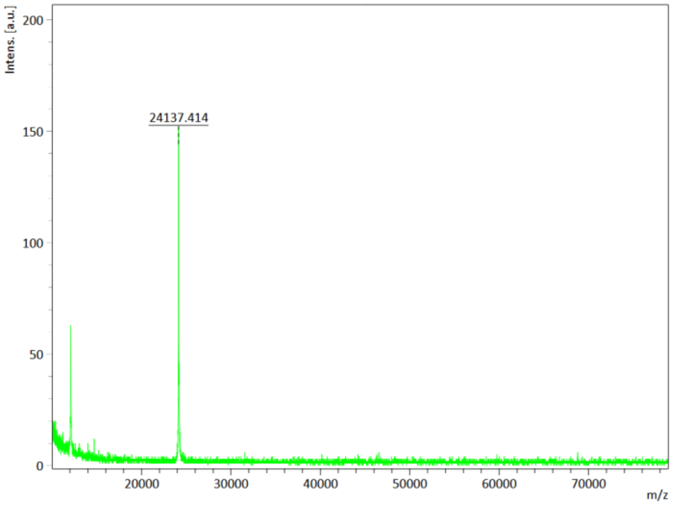
Intens. [a.u.]
200
150
100
50
0
24137.414
20000
30000
40000
50000
60000
70000
m/z
